# Mitochondrial temperature homeostasis resists external metabolic stresses

**DOI:** 10.1101/2023.05.24.542069

**Authors:** Mügen Terzioglu, Kristo Veeroja, Toni Montonen, Teemu O. Ihalainen, Tiina S. Salminen, Paule Bénit, Pierre Rustin, Young-Tae Chang, Takeharu Nagai, Howard T. Jacobs

**Author notes:** Corresponding author: Howard T. Jacobs, Faculty of Medicine and Health Technology, FI-33014 Tampere University, Finland. who shares corresponding and senior authorship.

## Abstract

Based on studies with a fluorescent reporter dye, Mito Thermo Yellow, and the genetically encoded gTEMP ratiometric fluorescent temperature indicator targeted to mitochondria, the temperature of active mitochondria in four mammalian and one insect cell-line was estimated to be up to 15 °C above that of the external environment to which the cells were exposed. High mitochondrial temperature was maintained in the face of a variety of metabolic stresses, including substrate starvation or modification, decreased ATP demand due to inhibition of cytosolic protein synthesis, inhibition of the mitochondrial adenine nucleotide transporter and, if an auxiliary pathway for electron transfer was available via the alternative oxidase, even respiratory poisons acting downstream of OXPHOS complex I. We propose that the high temperature of active mitochondria is an inescapable consequence of the biochemistry of oxidative phosphorylation and is homeostatically maintained as a primary feature of mitochondrial metabolism.

**IMPACT STATEMENT:** Mitochondria are up to 15 °C hotter than their external environment in living cells. In response to diverse metabolic stresses, mitochondrial temperature re-adjusts to this value whenever possible.

## INTRODUCTION

The mitochondrial system of oxidative phosphorylation (OXPHOS) is considered the most efficient energy conversion platform of non-photosynthetic eukaryotes. However, it does not operate at or anywhere near 100% thermodynamic efficiency. Much of the free energy released by the reoxidation of primary electron carriers such as NADH is converted to heat. Note that this applies under normal physiological conditions^1–3^, not only in cells expressing uncoupler proteins or treated with a chemical uncoupler. Furthermore, this heat production is not mere ‘waste’, since it is a major and regulatable energy source for maintaining body temperature in homeotherms such as mammals and birds^4^, and possibly many other organisms, even those living at a wide range of temperatures in varying environments. Uncoupler proteins merely serve an auxiliary function under cold stress, as shown by the fact that, in mice, ablation of the major uncoupling protein UCP1 is nonlethal^5^ and does not affect basal body temperature.

Using the mitochondrially targeted fluorescent dye Mito Thermo Yellow (MTY), it was recently shown that intramitochondrial temperature in respiring, human HEK293-derived (immortalized embryonic kidney) cells is at least 10-12 °C higher than the ambient temperature at which the cells are maintained^6^. In essence, this inference was based on the fact that the MTY fluorescence change, after cells were treated with OXPHOS inhibitors such as rotenone (targeted on OXPHOS complex I, cI), antimycin (targeted on OXPHOS complex III, cIII), oligomycin (targeted on OXPHOS complex V, cV) or subjected to prolonged anaerobiosis indicated a temperature drop of this magnitude, based on internal calibration at the end of each experiment. Inferences from the use of protein-based fluorescence probes targeted to mitochondria are similar^7^. In addition, similar methods have indicated that the endoplasmic reticulum (ER)^8^, cell nucleus^9, 10^, plasma membrane^11^ and different cellular compartments such as the cell body and neurites of neurons^12^, are at temperatures distinct from the extracellular environment and from each other.

These findings have far-reaching implications for biochemistry and physiology, but need further validation and elaboration before they can be generally accepted. In particular, since the cells of many organisms, notably homeotherms, are unable to tolerate prolonged high or low temperatures when applied externally, the question arises as to how much the temperature of mitochondria is permitted to vary *in vivo*, and whether and how it is regulated according to physiological conditions^13^.

In this study we undertook to assess mitochondrial temperature in a number of cultured cell-lines from different organisms, and subjected to different metabolic stresses involving well-characterized inhibitors and the transgenic expression of the thermogenic alternative oxidase from a marine invertebrate. Broadly, the outcome of these studies verifies that mitochondrial temperature is physiologically maintained at ∼15 °C above ambient temperature, and is subject to homeostatic regulation in response to metabolic stress.

## RESULTS

### Based on MTY fluorescence mitochondrial temperature is up to ∼15 °C above ambient

In previous studies^6^, MTY fluorescence was tracked in continuously oxygenated human HEK293-derived cells maintained at a constant external temperature, in which mitochondria had been prelabelled with MTY and then treated with one of several OXPHOS inhibitors under controlled oxygen concentration. Internal calibration in each experiment indicated a mitochondrial temperature decrease of 10-12 °C (Ref. 6) following the transition from normoxia to anaerobiosis, or inhibition of the OXPHOS system by classic inhibitors (rotenone, antimycin, cyanide or oligomycin – see Fig. 1). We conducted a systematic follow-up study with different inhibitors and cell-lines, and sufficient repeats to generate statistical reliability (Fig. 2), once the MTY fluorescence intensity had reached a stable value following initial oxygenation. See Figs. S1 and S2 for details of the instrumentation, its validation and calibration, and various steps that we undertook to validate the method, showing that the fluorescence signal of MTY in cell-free solution decreases almost linearly with temperature (Fig. S2D), is not responsive to changes in calcium concentration (Fig. S2E), hydrogen peroxide (Fig. S2F), or pH (Fig. S2G), across the physiological range of these parameters. Note that the curve generated (Fig. S2D), cannot be used directly for calibration in living cells, due to the endogenous contributions of autofluorescence and fluorescence quenching, which are considerable, and will vary according to how much label is actually taken up and retained in mitochondria in each trial. Instead, the internal calibration carried out in each experiment provides a more reliable guide to the temperature changes produced.

**Figure 1.**
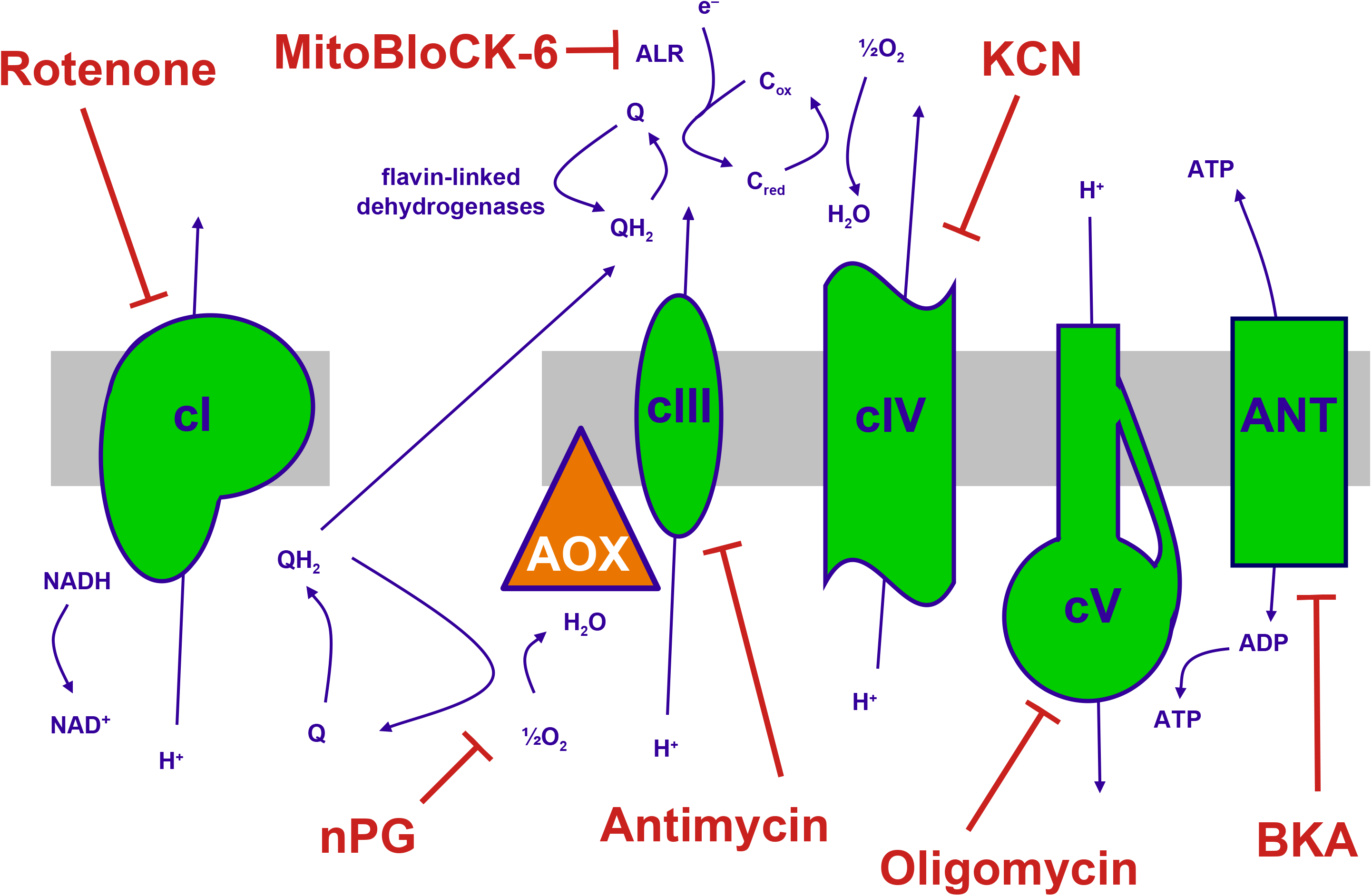
The mitochondrial OXPHOS system and inhibitors. Summary of the major components of the OXPHOS system and classic inhibitors. Protonmotive OXPHOS enzyme complexes (cI, cIII, cIV, cV) shown in green, the non-protonmotive transgenically introduced alternative oxidase (AOX) from *C. intestinalis* in orange, the inner mitochondrial membrane in grey. Ions and small molecules indicated in purple, inhibitors in brick-red. BKA – bongkrekic acid, an inhibitor of the adenine nucleotide translocase (ANT). nPG – *n*-propyl gallate, an inhibitor of AOX, Q – ubiquinone (oxidized coenzyme Q), QH_2_ – ubiquinol (reduced coenzyme Q), c_red_, c_ox_ – reduced and oxidized forms of cytochrome *c*, respectively. Complex I (cI), along with a number of flavin-linked dehydrogenases, one of which is succinate dehydrogenase, also described as complex II, each reduce ubiquinone, thus contributing input electrons to complex III (cIII). ALR – ‘augmenter of liver regeneration’, Evr1 in yeast, acts as an additional feeder of electrons to cytochrome *c*, from the oxidation of sulfhydryl groups in proteins destined for the mitochondrial intermembrane space, in the Mia40 pathway and is inhibited by MitoBLloCK-6.

**Figure 2.**
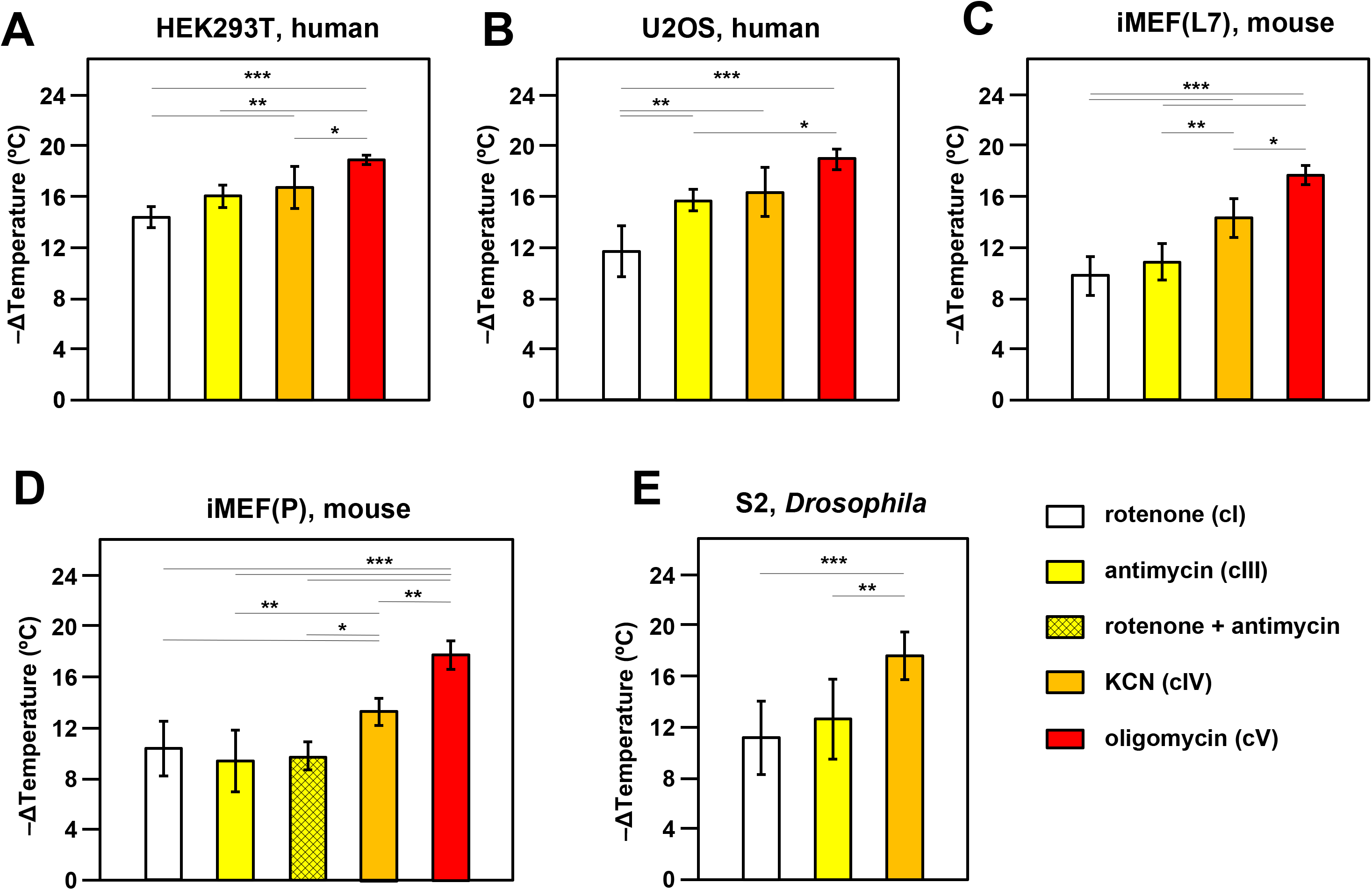
OXPHOS inhibitors decrease mitochondrial temperature to different extents. Extrapolated mitochondrial temperature shifts (means ± SD for at least 5 independent experiments in each case), based on MTY fluorescence in different cell-lines treated with the indicated OXPHOS inhibitors, with internal temperature calibration at the end of each experiment. *, **, *** denote significantly different data sets (one-way ANOVA with Tukey HSD *post-hoc* test, p< 0.05, 0.01 and 0.001, respectively). Note that a stable reading could not be obtained for S2 cells treated with oligomycin (Fig. S3B), hence reliable calibration was not possible. iMEF(L7) and iMEF(P) are two isolates of immortalized, embryonic fibroblasts derived from wild-type C57Bl/6J mice.

In most cell-lines tested there was a clear gradation of mitochondrial temperature decrease using inhibitors that act further along the OXPHOS chain: the effect was always greatest after oligomycin (inhibitor of ATP synthase or OXPHOS complex V, cV) and generally least for rotenone (cI inhibitor), with potassium cyanide (inhibitor of OXPHOS complex IV, cIV) and antimycin (cIII inhibitor) intermediate (Fig. 2). In general, findings were similar for different cell-lines, with an implied temperature decrease of the order of 17-19 °C produced by oligomycin. This implies a mitochondrial temperature of at least 54 °C, assuming that cV inhibition abolishes all mitochondrial heat production, or at least 52 °C, taking account of the possible slight deviation from linearity seen in the MTY fluorescence temperature-response curve at higher temperatures (dotted line, Fig. S2D). Two different isolates of immortalized mouse embryonic fibroblasts (iMEFs) gave similar results, with no significant difference in the extent of mitochondrial temperature decrease produced by rotenone to inhibit cI, by antimycin to inhibit cIII (Fig. 2C. 2D) or a combination of the two drugs (Fig. 2D). Unlike rotenone plus antimycin (Fig. S3A, panel i) other combinations of OXPHOS inhibitors did not give stable fluorescence values (Fig. S3A, panels ii-vi) and were not considered further. We were also unable to estimate directly the individual contribution to mitochondrial temperature maintenance of the ALR (Evr1) pathway (of mitochondrial protein import to the inter-membrane space), in which electrons are fed directly to cytochrome *c* (Fig.1): preparations of the ALR inhibitor MitoBloCK-6 (Ref. 14) that we were able to source had a distinct yellow colour that rendered measurements of MTY fluorescence uninterpretable (Fig. S3B).

We investigated the effect of culturing iMEF cells in low-glucose medium, with or without galactose supplementation, aiming to enhance dependence on mitochondrial respiration^15^. This culture-medium alteration had no significant effect on the effect of cI inhibition on mitochondrial temperature (Fig. S3C).

The inferred temperature of mitochondria in S2 cells from the fruit-fly *Drosophila*, a poikilotherm, when suspended and oxygenated at 25 °C, showed similar responses to OXPHOS inhibition as mammalian cells suspended at 37 °C, except that we were unable to derive a stable reading after oligomycin treatment (see Fig. S3D, panel i). MTY fluorescence was still increasing 2 hours after the addition of oligomycin, whereas other inhibitors, such as KCN, when applied to S2 cells, gave a stable reading over the same time period (e.g. Fig. S3D panel ii). Fluorescence reached a stable value after the addition of oligomycin in all of the other cell-lines tested, e.g. U2OS human osteosarcoma cells as shown (Fig. S3D panel iii).

### Protein ratiometric fluorescence confirms mitochondrial temperature measured by MTY

Although the previous study excluded the most obvious possible artefacts from the use of MTY fluorescence as a mitochondrial thermometer^6^, such as an effect of membrane potential, the redox state of the respiratory chain or reactive oxygen species (ROS) production, artefacts of unknown provenance could always be in play. We therefore set about applying a different method to measure mitochondrial temperature in these cell lines, based on the ratiometric fluorescence of mito-gTEMP^7^. This technique is based on the fluorescence ratio between two previously developed fluorescent proteins, mT-Sapphire and Sirius, co-expressed as a single, self-cleaved polypeptide, with each cleavage product targeted to mitochondria using a reiterated cytochrome oxidase subunit 8 (COX8) presequence (Fig. 3A). The basis of the method is the fact that the fluorescence of Sirius is highly temperature sensitive, declining (like MTY) as temperature rises, whereas that of mT-Sapphire is not^7^. We stably transfected HEK293T, U2OS and iMEF(P) cells with mito-gTEMP and confirmed mitochondrial targeting, as exemplified in Fig. 3B and Fig. S4A. Since the two proteins are expressed at a fixed stoichiometry, the peak fluorescence ratio is independent of expression level, which may vary even within a cloned mammalian cell population^16^, as seen, for example, in Fig. 3B.

**Figure 3.**
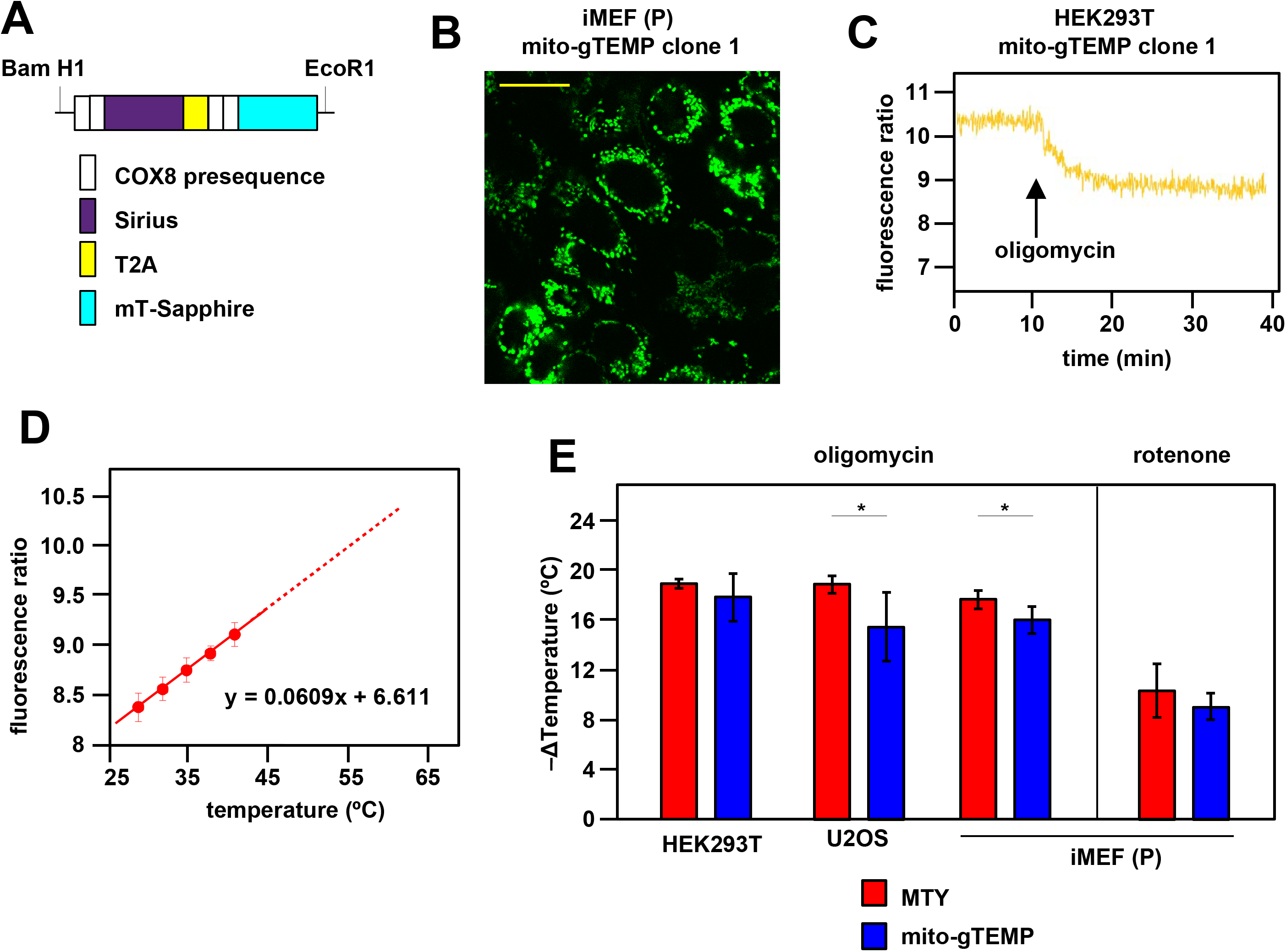
Ratiometric fluorescence probes confirm mitochondrial temperature measurements by MTY. (A) Diagrammatic map (not to scale) of the self-cleaving polyprotein encoded by plasmid mito-gTEMP (coxVIIIx2-gTEMP)/pcDNA3, indicating the reiterated COX8 targeting presequences, the Sirius and mT-Sapphire coding regions and the T2A element^76^, which promotes self-cleavage via ribosome frameshifting^77^. (B) Illustrative mT-Sapphire fluorescence image of iMEF cell clone expressing mito-gTEMP, showing variable expression even within cells of the clone. Scale bar 5 μm. See Fig. S4A for confirmation of mitochondrial localization. (C) Illustrative trace of mT-Sapphire/Sirius fluorescence ratio in HEK293T cell clone expressing mito-gTEMP, following treatment with oligomycin. Note that, as for MTY, Sirius fluorescence decreases with increasing temperature, so that the decreased mT-Sapphire/Sirius fluorescence ratio after oligomycin treatment represents a decrease in temperature. (D) Calibration curve for mT-Sapphire/Sirius fluorescence ratio (means ± SD, 9 independent experiments in each case) in mito-gTEMP expressing iMEFs treated with oligomycin, and shifted to various temperatures up to 42 °C. Solid red line represents linear best fit (as per equation shown) extrapolated to higher temperatures (dotted red line) that were not directly tested due to the effects of prolonged high temperatures on cell integrity and viability^17^. Note that, during these trials, we observed that continuous illumination of mito-gTEMP expressing cells at the excitation wavelength for more than 40 min caused damage affecting the fluorescence signal. All calibration measurements were therefore conducted over < 35 min. We also noted that the internal temperature of the cuvette took ∼2 minutes to equilibrate with that of the surrounding Peltier (Fig. S2), for which reason fluorescence values were measured over just the final minute at each calibration step. (E) Extrapolated mitochondrial temperature shifts (means ± SD for at least 5 independent experiments in each case) in the indicated cell-lines and mito-gTEMP transfected cell clones following treatment with oligomycin, based on on (red bars) MTY fluorescence, reproduced from corresponding panels of Fig. 2, and (blue bars) mito-gTEMP fluorescence ratio, applying calibration curves such as shown in panel (C), derived separately for each cell line studied. ** denotes significant differences between the two methods (Student’s t test within each cell-line, p< 0.01).

To check for possible artefacts introduced by mito-gTEMP in the presence of OXPHOS inhibitors we checked whether the drugs alone in PBS affect background fluorescence in the absence of cells. Background fluorescence at the wavelengths used to track mT-Sapphire and Sirius were both more than an order of magnitude less than that of the fluorescent proteins expressed in iMEFs, and were unaffected by rotenone (Fig. S4B). Background fluorescence at both wavelengths was slightly increased by oligomycin (Fig. S4C), but the fluorescence ratio was unchanged (Fig. S4E). In contrast, antimycin produced a substantial change in background fluorescence at the Sirius wavelength (Fig. S4D), which affected the fluorescence ratio (Fig. S4F), making it unsuitable for use in estimating mitochondrial temperature shifts.

Since, based on MTY fluorescence, we had inferred a maximal amount of mitochondrial temperature decrease in the presence of oligomycin, we used oligomycin-treated cells to calibrate the mito-gTEMP fluorescence ratio against externally imposed temperature. This assumes that any residual mitochondrial heat production in oligomycin-treated cells is negligible and invariant with respect to externally applied temperature changes.

After oligomycin treatment, iMEF mitochondria showed a decline in the mito-gTEMP fluorescence ratio over ∼10 min, thereafter holding a constant value for at least a further 25 min that is assumed to represent the ambient temperature (Fig. 3C). This enabled us to construct an *in vivo* calibration curve for the mito-gTEMP fluorescence ratio against externally imposed temperature (Fig. 3D), which fitted a linear function over the temperature range at which we were able to conduct the measurement: higher temperatures result in loss of cell integrity and viability^17^, so were not used. Extrapolation to higher mitochondrial temperatures is therefore based on the assumption that the relationship remains linear, as was determined previously for the isolated gTEMP proteins^7^.

Oligomycin treatment of mito-gTEMP expressing cell clones derived from three different mammalian cell-lines gave extrapolated values for the mitochondrial temperature decrease in the same range as those inferred by MTY, although slightly (1-2 °C) less in each case, implying an internal mitochondrial matrix temperature of ∼52-54 °C (Fig. 3E). Rotenone treatment of iMEF(P) cells also produced a similar amount of mitochondrial temperature decrease when estimated by mito-gTEMP as by MTY (Fig. 3E).

The slight discrepancy in the mitochondrial temperature inferred by the two methods prompted us to consider whether mito-gTEMP and MTY are targeted to different sub-mitochondrial compartments. The matrix localization of GFP-related fluorescent reporters using the COX8 targeting peptide, is well documented^18^. However, the exact intramitochondrial location of MTY is not known, besides the fact that at least some of it interacts with the matrix protein aldehyde dehydrogenase 2, ALDH2 (Ref. 19). To address this issue experimentally, we sub-fractionated mitochondria from cells pre-stained with MTY (see Fig. S2H) and measured fluorescence at 562 nm, subtracting the autofluorescence manifested by the corresponding fraction from unstained cells prepared in parallel. To prevent dye leakage, we supplied a standard pyruvate-glutamate-malate substrate mix to maintain mitochondria in the energized state throughout the procedure. In the first fractionation step, mitoplasts were separated from the outer membrane/inter-membrane space fraction. In two trials, the mean proportion of MTY fluorescence retained in mitoplasts was 92%. In the second fractionation step, mitoplasts were sonicated to prepare fractions enriched for matrix versus ‘inside-out’ sub-mitochondrial particles (SMPs). In two trials, the mean proportion of MTY fluorescence retained in the SMP fraction was 90%. These findings are consistent with MTY being retained within, or closely associated with, the inner mitochondrial membrane.

### Individual cells and organelles show mitochondrial temperature differences and fluctuations

As studied by either method, mitochondrial temperature represents an average over many cells, each containing many mitochondria. We used confocal imagining to look at the profile of mitochondrial temperature in individual iMEF(P) cells and organelles stained with MTY (Fig. 4A, panel i). This revealed much greater heterogeneity between and even within cells not seen, for example, with the ubiquitous mitochondrial stain MitoTracker^TM^ Deep Red FM (Fig. 4A, panel ii). Within individual cells stained with MTY we observed some more brightly staining puncta (Fig. 4B; Supplementary Movie S1). Time-lapse imaging revealed that mitochondria in individual cells were cooled to variable extents by oligomycin treatment (Fig. 4C).

**Figure 4.**
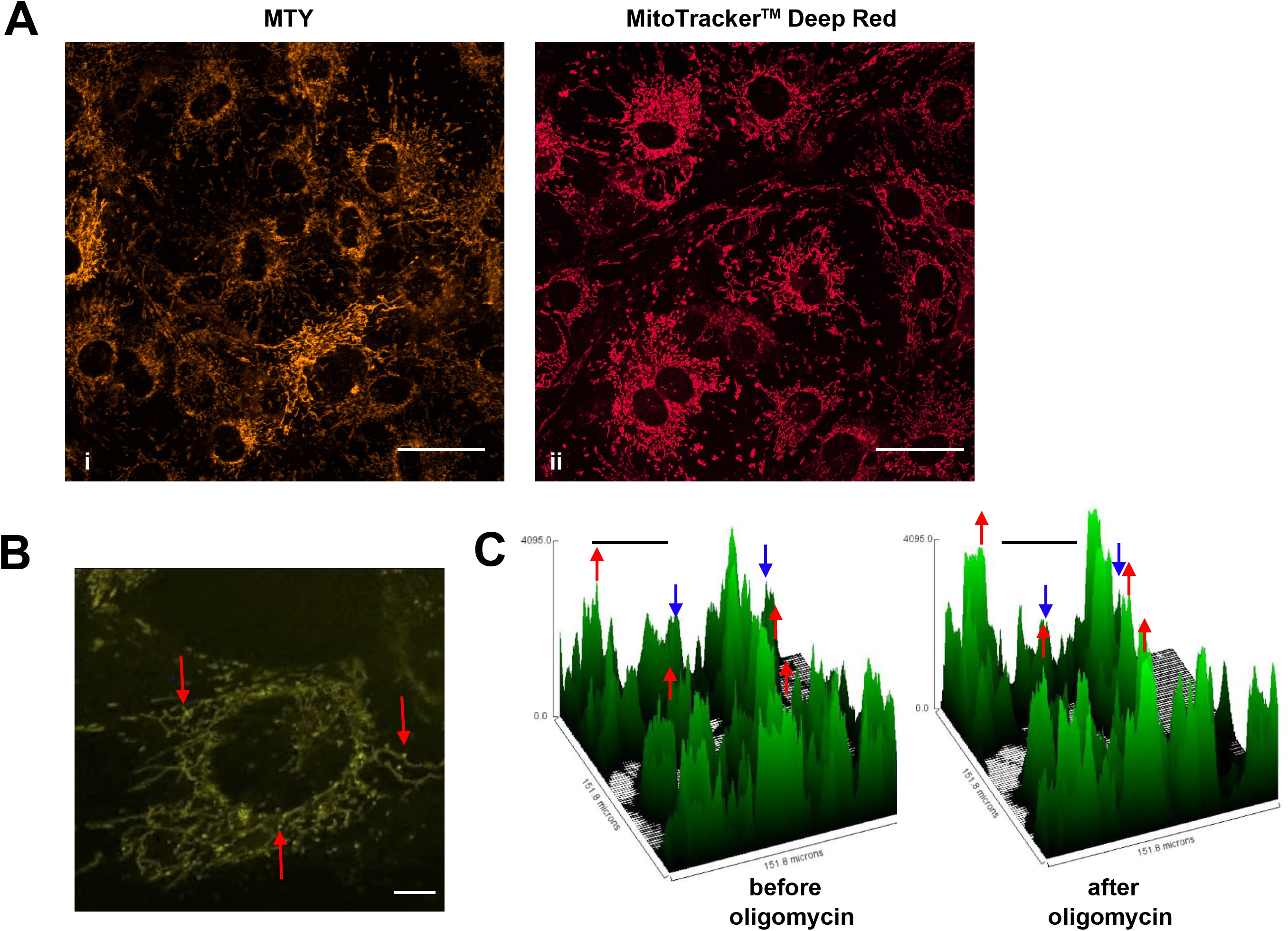
Mitochondrial temperature variation between and within cells. (A, B) Micrographic images of iMEF cells stained with MTY (A, panel i and B), alongside a parallel culture of cells stained with Mitotracker^TM^ Deep Red FM (A, panel ii), which shows much more uniform staining. The intensity of MTY staining appears to vary between and even within cells. Scale bars – 10 μm. (B) Still image from Supplementary Movie S1, brightness and contrast optimized, showing bright (i.e., potentially cooler) foci (arrowed) within the less intensely stained mitochondrial network. Many of these foci appear mobile within the mitochondrial network, over the 5 min time-scale of the movie. Scale bar – 2 μm. (C) Fluorescence Intensity histograms of a field of cells immediately before and 33 min after oligomycin treatment. Note that, in addition to the general pattern of increased brightness (e.g., peaks represented by red dots), mitochondria in some cells showed only minor changes in fluorescence intensity (e.g., peaks denoted by blue dots). Scale bars – 50 μm.

### Metabolic adaptation can maintain or restore mitochondrial temperature

When mito-gTEMP expressing iMEF cells were incubated in PBS at 37 °C without the addition of any inhibitor we observed that, after reaching a steady fluorescence ratio indicative of a temperature of 53-54 °C, there was a modest but progressive decline in the fluorescence ratio, such that by 35 min of incubation the inferred mitochondrial temperature was around 4 °C lower than the peak value reached at approximately 10 min (Fig. 5A, 5B). Although remaining much warmer than the surrounding medium, this suggests that, in the complete absence of an externally supplied, metabolizable substrate, mitochondrial metabolism adjusts to a slightly lower heat output and steady-state temperature. To explore this further we added the main substrates normally provided to mammalian cells in culture, namely glucose and glutamine. As expected, this enabled mitochondria to maintain a higher temperature of ∼53 °C at the end of the experiment, although this was still slightly less than the initial temperature peak of ∼55 °C between 5-10 min of incubation (Fig. 5B, 5C). However, the addition of glucose and glutamine at 10 min did produce a small rise in mitochondrial temperature (Fig. 5B, 5D). In sum, these data indicate that mitochondrial temperature fluctuates within a narrow range according to substrate supply.

**Figure 5.**
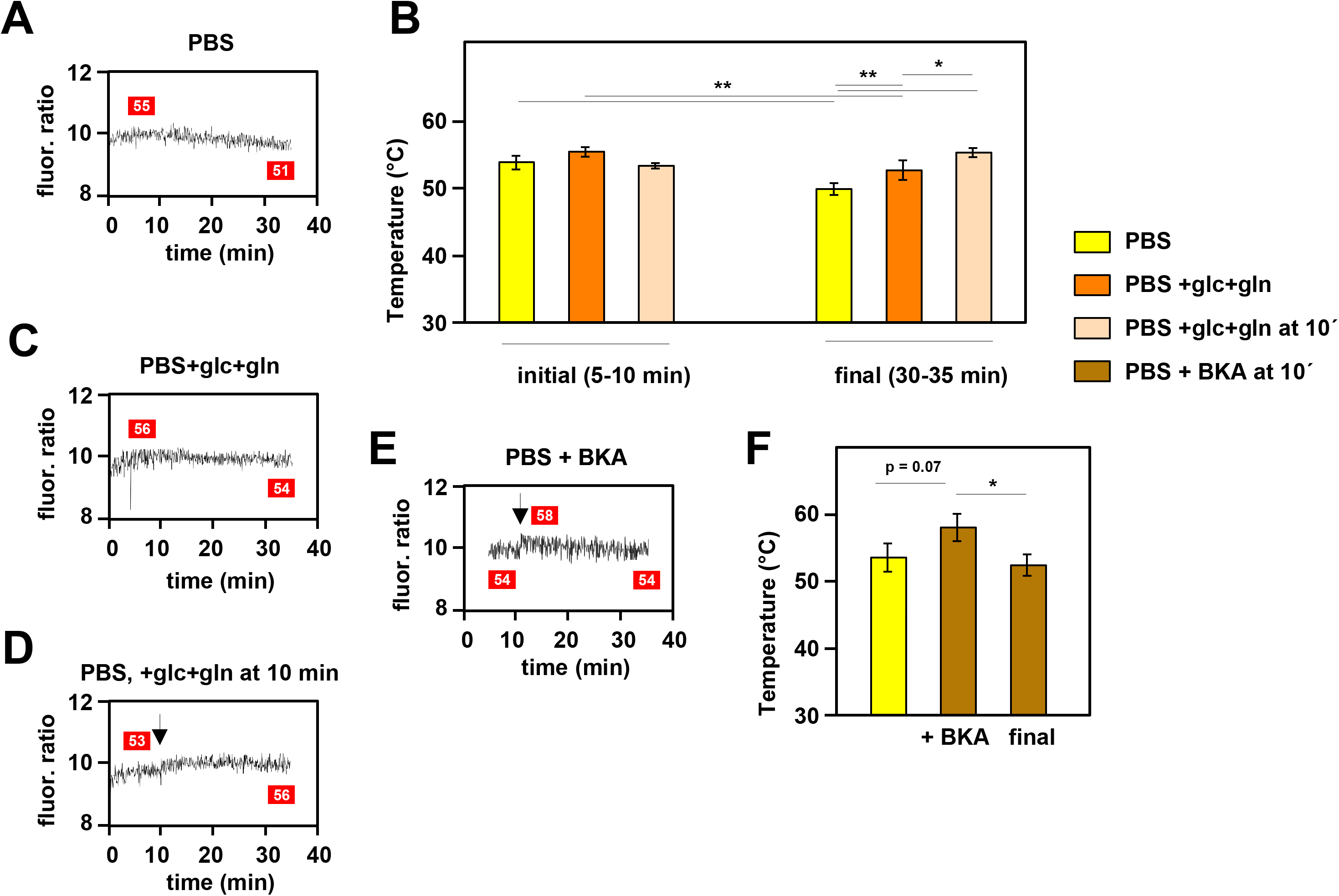
Mitochondrial temperature fluctuates in response to substrate availability. (A, C, D, E) Representative fluorescence ratio traces of mito-gTEMP expressing iMEF(P) cells resuspended in various media without OXPHOS inhibitors. PBS – phosphate-buffered saline, glc – glucose and gln – glutamine at standard cell-culture concentrations, BKA – bongkrekic acid. Arrows indicate time of addition of substrates or of BKA. Mitochondrial temperatures (°C) extrapolated from calibration curve (Fig. 3D) shown in red boxes. (B) Mitochondrial temperatures (°C, means ± SD of 5 independent experiments), averaged over the time intervals since cell resuspension as shown, extrapolated from mito-gTEMP fluorescence ratios using calibration curve (Fig. 3D), for cells with and without substrate additions as shown. *, **, *** denote statistical significance (p < 0.05, 0.01, 0.001, respectively, one-way ANOVA with Tukey *post-hoc* HSD test). (F) Mitochondrial temperatures (°C, means ± SD, n = 3 independent experiments), averaged over the time intervals since cell resuspension as shown, extrapolated from mito-gTEMP fluorescence ratios using calibration curve (Fig. 3D), for cells treated with BKA at 11 min (values averaged over 11-16 min) and then tracked until 35 min (final values averaged over 30-35 min). *, **, *** denote statistical significance (p < 0.05, 0.01, 0.001, respectively, one-way ANOVA with Tukey *post-hoc* HSD test).

In order to explore the issue further, we treated cells with bongkrekic acid (BKA), a potent inhibitor of the mitochondrial adenine nucleotide translocase, ANT (Ref. 20). The ANT exchanges ADP and ATP across the mitochondrial inner membrane (IMM), thus enabling OXPHOS to supply the rest of the cell with ATP. Previous investigators have shown that cultured mouse fibroblasts respond to BKA by inducing glycolysis and downregulating mitochondrial respiration, whilst maintaining a high mitochondrial membrane potential^21^. In initial trials we found that BKA quenches the fluorescence of MTY, even in the absence of cells (Fig. S4G). Therefore, we used the mito-gTEMP fluorescence ratio, which was less disturbed by the drug, to estimate its effects on mitochondrial temperature. We reasoned that the metabolic switching induced by BKA should result in a substantial mitochondrial temperature decrease, due to the expected lowering of flux through the respiratory chain. However, the mitochondrial temperature decrease that we observed was minimal, following an apparent slight increase upon addition of the drug (Fig. 5E, F), qualitatively consistent with the MTY signal after discounting the quenching effect (Fig. S4G).

### AOX maintains or restores high mitochondrial temperature in the presence of OXPHOS inhibitors

Next we looked at iMEF cells constitutively expressing the non-protonmotive alternative oxidase (AOX) from *Ciona intestinalis*, which provides an alternative pathway for respiratory electron flow from ubiquinol to oxygen, if cIII and/or cIV are inhibited^22^. We first verified by respirometry that iMEF AOX cells were resistant to inhibitors of cIII (antimycin, Fig. S5A) and cIV (KCN, Fig. S5B).

In cells that principally metabolize cI-linked substrates, AOX makes only a minor contribution to respiratory electron flow under non-inhibited conditions^23^. However, if cI is blocked, forcing cells to switch to other ubiquinone-linked substrates, AOX becomes active^23^. Once again, the cell must switch substantially to glycolysis to sustain ATP levels, and to cytosolic NADH reoxidation via lactate dehydrogenase.

Consistent with these expectations, the initial mitochondrial temperature decrease produced by the cI-inhibitor rotenone in AOX-expressing iMEFs was very similar to that of control iMEFs, with a mitochondrial temperature decrease of ∼10 °C based both on MTY fluorescence (Fig. 6A, 6B) and the mito-gTEMP fluorescence ratio measured in iMEF cells stably transfected with AOX (Fig. 6B, Fig. S6A). However, this was followed by a gradual mitochondrial rewarming (Fig. 6A panel i, 6B, 6C). AOX-expressing iMEFs treated with other OXPHOS inhibitors showed a similar behaviour (Fig. 6A panels ii-iv, 6C), except that the initial mitochondrial temperature decrease was substantially less than that seen in control iMEFs (Fig. 6A panels ii-iv, 6B), whilst the rate of rewarming differed according to the inhibitor used (Fig. 6C). We were not able to estimate directly the contribution of AOX to temperature maintenance, due to fluorescence quenching by the AOX inhibitor n-propyl gallate, nPG (Fig. S6B).

**Figure 6.**
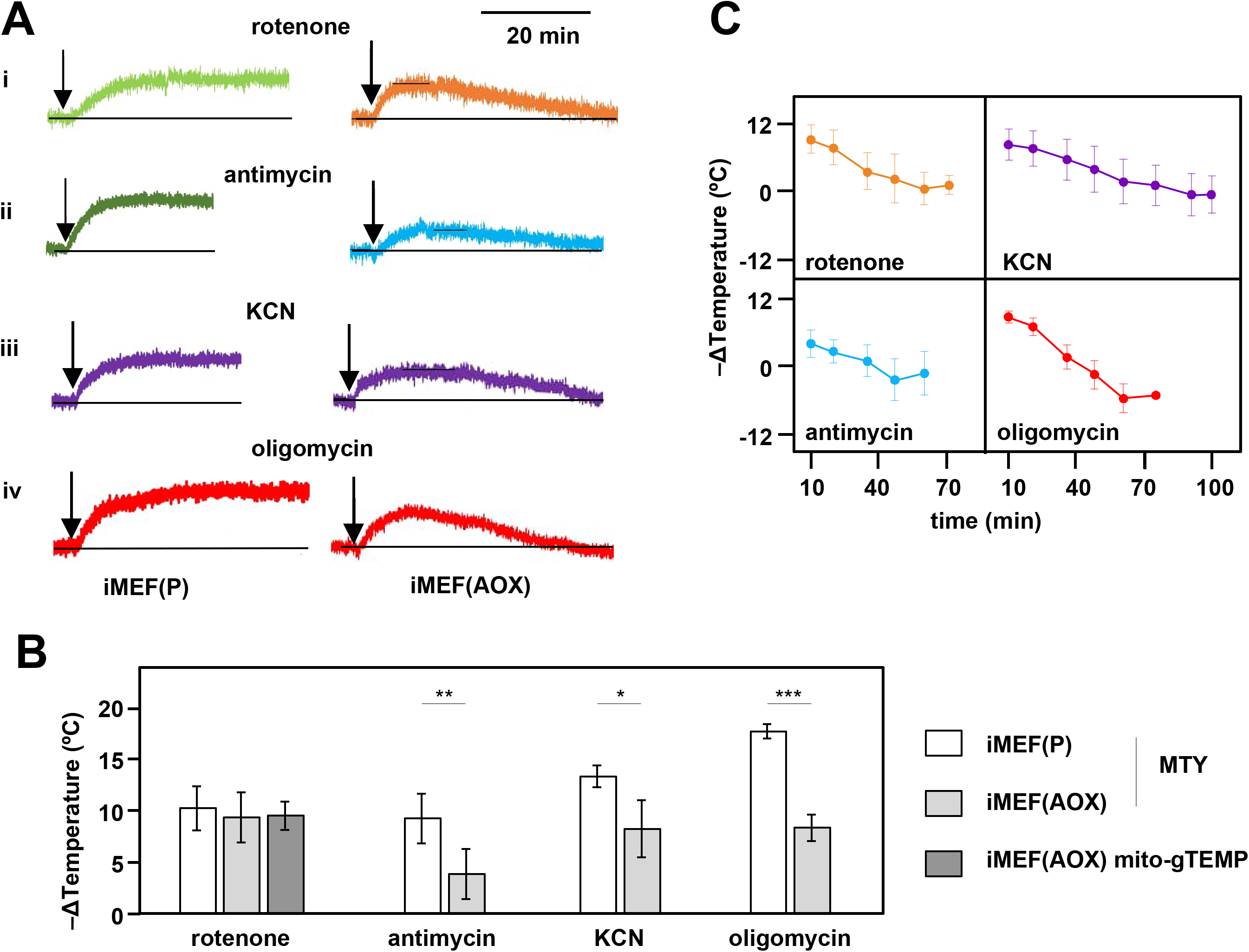
Restoration of high mitochondrial temperature in AOX-expressing cells after OXPHOS inhibition. (A) Representative traces of MTY fluorescence over time, in control and AOX-expressing iMEFs treated with the OXPHOS inhibitors shown. Vertical axes are arbitrary values, horizontal axes scaled as indicated. Note that increased fluorescence indicates cooler mitochondria. Control iMEF(P) cells reached and maintained plateau levels of fluorescence, indicating a sustained decrease in mitochondrial temperature, whilst iMEF(AOX) cells showed rewarming after reaching a peak fluorescence value. (B) Peak mitochondrial temperature decrease (means ± SD, n ≥ 4 independent experiments in the indicated cell-lines following treatment with OXPHOS inhibitors, inferred from MTY fluorescence or mito-gTEMP fluorescence ratio as shown. Horizontal bars indicate significant differences for pairwise comparisons between iMEF(P) and iMEF(AOX) cells treated with the indicated inhibitors (MTY data, Student’s t test, p < 0.05, 0.01 and 0.001, denoted, respectively, by *, ** and ***). There was no significant difference between the methods, applied to rotenone-treated iMEF(AOX) cells (Student’s t test, p > 0.05). (C) Mitochondrial rewarming (means ± SD, n ≥ 5 independent experiments except at latest time-points, where n = 3 or 4) in iMEF(AOX) cells treated with the inhibitors shown, inferred from MTY fluorescence traces.

### Mitochondrial temperature is maintained despite varying ATP demand

To model the effects on mitochondrial temperature of a more physiologically relevant perturbation, we downregulated cytosolic protein synthesis, the major cellular consumer of ATP, by treating iMEFs with anisomycin (Fig. 7). Using our earlier calibrations, we initially estimated the mitochondrial temperature in cells prior to anisomycin treatment, then followed the effect of anisomycin treatment for up to 18 hours (Fig. 7). In contrast to our expectation that anisomycin, by attenuating ATP demand, would decrease electron flux in the OXPHOS system and thus lead to a drop in mitochondrial temperature, the latter instead showed a transient slight increase (Fig. 7A).

**Figure 7.**
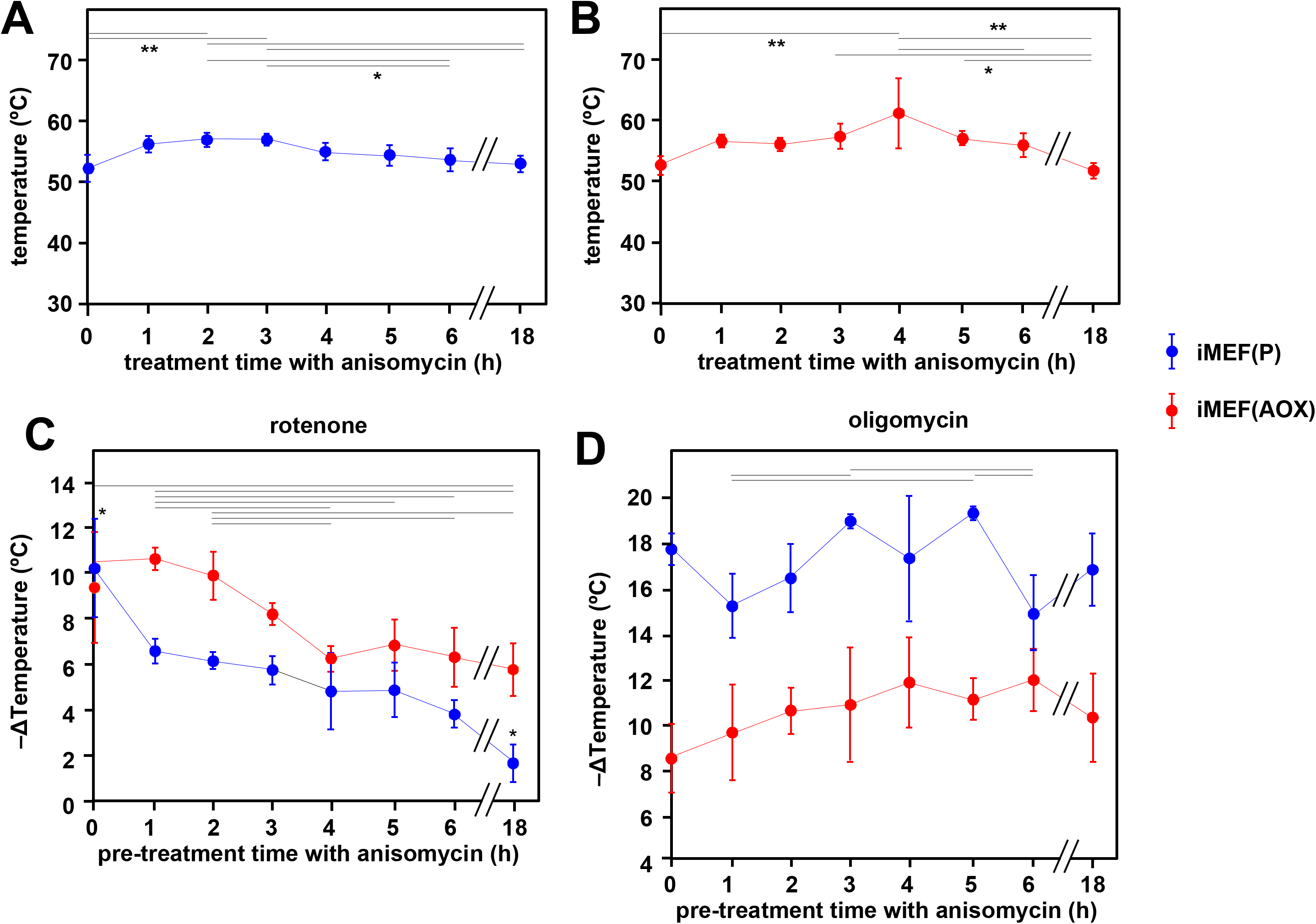
Cells maintain high mitochondrial temperature despite varying ATP demand. (A, B) Intramitochondrial temperatures (means ± SD) estimated by mito-gTEMP fluorescence ratio in iMEF(P) and iMEF(AOX) cells as indicated, treated with 150 μM anisomycin for the times shown. Temperatures extrapolated from calibration curve shown in Fig. 3D. Horizontal bars denote data points that were significantly different (one-way ANOVA with Tukey HSD test, p < 0.05). (C) Extrapolated mitochondrial temperature shifts (means ± SD for at least 4 independent experiments in each case) based on MTY fluorescence, as indicated, in iMEF(P) and iMEF(AOX) cells pre-treated with 150 μM anisomycin for the times shown, followed by treatment with rotenone. Zero-time values (i.e. cells not treated with anisomycin) are reproduced from Figs. 2D and 5B for iMEF(P) and iMEF(AOX) cells, respectively. Values for the two cell lines are significantly different at each time-point except 0 and 4 h (Student’s t test, p < 0.05). Within each cell-line, in addition to the clear downward trend with time (i.e., lesser amount of mitochondrial temperature decrease), the values for the extreme time points (0 and 18 h) are significantly different from each other and from all other time-points for the iMEF(P) line (denoted by asterisks), whilst for the iMEF(AOX) line significant pairwise differences are indicated by horizontal lines above the graphed values (one-way ANOVA with Tukey *post-hoc* HSD test, p > 0.05). (D) Extrapolated mitochondrial temperature shifts (means ± SD for at least 4 independent experiments in each case) based on MTY fluorescence, as indicated, in iMEF(P) and iMEF(AOX) cells pre-treated with 150 μM anisomycin for the times shown, followed by treatment with oligomycin. Zero-time values (i.e. cells not treated with anisomycin) are reproduced from Figs. 2D and 5B for iMEF(P) and iMEF(AOX) cells, respectively. Values for the two cell lines are significantly different at each time-point (Student’s t test, p < 0.05). Within each cell-line, and despite the apparent trend of the means for iMEF(AOX) cells, the values for different time points are, in general, not significantly different from each other (one-way ANOVA with Tukey *post-hoc* HSD test, p > 0.05). Exceptions for iMEF(P) cells are indicated by horizontal lines (p < 0.01).

Thereafter, high mitochondrial temperature was sustained for many hours, although by 18 hours of exposure to the drug many of the cells were no longer attached and probably dying (the cells analyzed in Fig. 7A were those that had remained attached). AOX-expressing iMEF cells (Fig. 7B) showed an almost identical behaviour, except that the cells remained healthy-looking even after 18 hours in anisomycin. To evaluate the contribution of cI and cV to the maintenance of mitochondrial temperature in anisomycin-treated cells, we stained anisomycin-pretreated iMEFs with MTY, then tracked fluorescence changes after treatment with rotenone (Fig. 7C) or oligomycin (Fig. 7D). The mitochondrial temperature of anisomycin-treated iMEFs showed a lesser effect of rotenone than iMEFs not treated with anisomycin (Fig. 7C, blue trace, compare 1-18 h time-points with 0 h control), and this effect was increasingly pronounced after longer times of anisomycin exposure.

Mitochondrial temperature in the surviving cells after 18 h in anisomycin was almost unaffected by rotenone. In contrast, and despite some experiment-to-experiment variation, anisomycin-exposed iMEFs were as affected by oligomycin as prior to treatment, even up to 18 h (Fig. 7D, blue trace).

AOX-expressing iMEFs behaved differently. After anisomycin treatment, they showed a consistently greater mitochondrial temperature decrease produced by rotenone than did control iMEFs not expressing AOX (Fig. 7C, red trace). Although the effect of oligomycin declined with increasing times of anisomycin exposure, it did so more gradually than for control iMEFs not expressing AOX. The mitochondria of anisomycin-treated iMEF(AOX) cells showed a temperature decrease by oligomycin significantly less than anisomycin-treated control iMEF(P) cells, regardless of the length of time of anisomycin exposure, but also showed a possibly increased effect of oligomycin, with increasing anisomycin exposure time (Fig. 7D, red trace), at least for the first hours. Anisomycin-exposed iMEF(AOX) cells showed the same slow rewarming after rotenone treatment (Fig. S7A panel i, ii) as did iMEF(AOX) cells not exposed to anisomycin (Fig. 5A panel i, 5C), whilst control iMEFs did not show any such effect from rotenone (Fig. S7A, panel iii, iv).

Conversely, the rewarming effect after oligomycin treatment was slower and of much lower magnitude for anisomycin-treated than untreated iMEF(AOX) cells (Fig. S7B panels i, ii), whilst a modest rewarming was also seen for control iMEFs not expressing AOX (Fig. S7B panel iii, iv).

## DISCUSSION

In this study, using the mitochondrially targeted, temperature-sensitive dye MTY applied to five cell-lines, we confirmed that physiologically active mitochondria *in vivo* exhibit high temperature, at least 15 °C above that of the external environment, i. e., ∼52 °C in mammalian cells. We validated the measurements using the mito-gTEMP system, based on the ratiometric fluorescence of two mitochondrially localized reporter proteins. We visualized mitochondrial temperature fluctuations at the single-cell level, and documented the maintenance of high mitochondrial temperature despite variations in growth conditions, substrate availability, ATP demand, and even OXPHOS poisons, if an alternative respiratory pathway is available.

No chemical reaction is 100% thermally efficient, and those of OXPHOS are no exception, with up to 60% of the energy yield from the biological oxidations of respiration and ATP synthesis released as heat^2, 3^. It has long been assumed that this heat was simply radiated to the rest of the cell or the external environment, maintaining intracellular temperature at a uniform 37-42 °C in mammals and birds, or at a variable, environmentally determined temperature in poikilotherms such as insects, generally considered as ectothermic. Our findings are inconsistent with this model, indicating instead that much of the heat generated by OXPHOS is retained within the organelle, keeping mitochondrial temperature far above that of other cell components and, whenever possible, within a narrow range. Heat conductance from the mitochondria might also be modulated to maintain mitochondrial temperature.

### Validation and calibration of high mitochondrial temperature

In previous studies using MTY in human HEK293 and HeLa cells we documented fluorescence changes after treatment with OXPHOS inhibitors that were indicative of high mitochondrial temperature^6, 24^. In the present study we extended these findings to other cell lines, including one from *Drosophila*, and two isolates of iMEFs, immortalized using a different cocktail of factors^25, 26^. We systematically conducted multiple repeats with each of five drugs impacting the OXPHOS system. In each case, we implemented an internal calibration step after MTY fluorescence had reached a plateau level in the presence of the inhibitor, by abruptly shifting the external temperature up, then down by 3 or 5 °C, at minimally 4 minute intervals. In each case, it is assumed that the observed fluorescence shift is due only to the change in the externally applied temperature (i.e., that any ‘corrective’ response of the cell in the presence of the inhibitor is negligible). This is supported by the fact that the new fluorescence values reached during the calibration steps are essentially constant over the time-scales used (see Fig. S2C for examples). Internal calibration in each experiment provides a control for variation in the exact number of cells or the amount of label taken up. The use of a Peltier jacket surrounding the cuvette in the apparatus ensures that temperature changes in the cuvette rapidly and accurately reflect those applied, as verified (Fig. S2A, S2B).

Importantly, the mito-gTEMP fluorescence ratio, and its calibration against externally applied temperature, provides a second check on the findings obtained using MTY. Although the principle of the mito-gTEMP method is different, being based on fluorescent reporter proteins rather than an externally applied dye, the calibration steps rely on the same, validated assumptions, and produced temperature readouts very similar to those derived using MTY, although typically several °C lower in the case of each of the mammalian cell-lines tested (Fig. 3E). As stated earlier, the MTY-based meaurements may be slight over-estimates (by ∼2 °C), due to the minor deviation from linearity of the MTY fluorescence temperature-response curve (Fig. S2D).

Although the two methods gave essentially concordant results, it is important to recognize that both have limitations. Apart from our failure to obtain a stable fluorescence reading for S2 cells treated with oligomycin, we did not experience problems working with MTY on any of the cell-lines tested in the presence of single OXPHOS inhibitors, such as were reported earlier for a human primary fibroblast line^24^. However, using combinations of OXPHOS inhibitors, other than rotenone plus antimycin, we were not able to infer mitochondrial temperature by the MTY method, due to unstable fluorescence readings (Fig. S3A). These are likely due to dye leakage resulting from membrane-potential anomalies of the type reported earlier^24^. In particular, we note that most combinations of OXPHOS inhibitors that failed to give a stable MTY fluorescence reading involved oligomycin plus an inhibitor of respiration. Since we already know that a complete loss of membrane potential leads to leakage of the dye, we surmise that collapse of membrane potential is the most likely reason for the fluorescence instability. Even in the presence of oligomycin, a minimal respiratory electron flow balanced against proton leakage should suffice to maintain a membrane potential. Similarly, even when respiration is inhibited, ATP synthase alone should be able to generate a membrane potential. However, the membrane potential may collapse when both oligomycin and a respiratory chain inhibitor are simultaneously applied.

Various confounding factors could potentially influence MTY fluorescence. In previous studies it was argued that drugs that have different effects on mitochondrial membrane potential nevertheless all lead to a sustained increase in MTY fluorescence. Since it is not possible to stain cells simultaneously with MTY and membrane-potential sensitive dyes such as TMRM, due to spectral overlap, we need to rely on experiments conducted under parallel conditions, both by ourselves^27^ and others^28, 29^. Treatment of cells with respiratory inhibitors such as rotenone or antimycin lead to substantially decreased membrane potential^27, 28^, whereas the effects of oligomycin on mitochondrial membrane potential are generally minor, and vary between cell-lines and over time. Commonly, oligomycin results in a small increase in mitochondrial membrane potential, sometimes followed by a decrease^28, 29^. In contrast, we here observed a sustained increased of MTY fluorescence in all of the mammalian cells treated with any of these inhibitors. This implies that any quantitative effect of membrane potential on MTY fluorescence is negligible, unless the potential is completely lost, leading to dye leakage. Any confounding effect upon MTY fluorescence of other physiologically varying parameters, such as calcium concentration, hydrogen peroxide or pH, was directly excluded (Fig. S2E, S2F, S2G) The excitation wavelength for mito-gTEMP (360 nm) falls within the ultra-violet range, and the resulting cellular damage under continuous illumination for more than 60 min renders calibration unreliable^30^. An alternative probe set, based on the recently developed B-gTEMP^25^, should overcome the problem in future studies. In the current report, we used mito-gTEMP only up to 35 min, which generated a reliable calibration curve (Fig. 3D). However, this is not long enough to profile very slow temperature changes such as the rewarming we observed using MTY on AOX-expressing cells (Fig. 6). In addition, we found that some inhibitors, notably antimycin (Fig. S4F), though not rotenone (Fig. S4B) or oligomycin (Fig. S4E), affected the mito-gTEMP fluorescence ratio, compromising its use. Similarly, the effects of BKA (Fig. S4G), nPG (Fig. S6B) and MitoBloCK-6 on MTY fluorescence readings preclude their use with the probe. One important caveat regarding the application of mito-gTEMP is that it was necessary to extrapolate the calibration curve beyond 42 °C, assuming the linearity seen earlier for the isolated proteins^7^. Above this temperature, *in vivo* calibration is not possible, due to the loss of cellular viability and integrity, accompanied by major changes in protein composition and distribution^17^.

The finding that mitochondrial temperature is of the order of 50 °C or more and distinct from that of the rest of the cell has been challenged on both theoretical^31^ and experimental grounds^32^. However, recent data indicate that the slow thermal diffusivity of mammalian cells^30^ is sufficient to account for the maintenance of intracellular temperature differences inferred under physiological conditions and further documented here. Furthermore, the maintenance of high mitochondrial temperature as a source of intracellular heat is supported by theoretical considerations^1^.

It should also be noted that previous studies using different probes have implied that the temperature across the cell is not uniform. Some organelles or active regions manifest distinct temperature inhomogeneities: notably, the nucleus is 1-2 °C warmer than the cytoplasm^9, 10^, the nucleolus even hotter^33^ and perinuclear mitochondria hotter than those at the cellular periphery^34^.

It will therefore be of great interest to use the temperature reporters employed here to track how the various treatments and stresses that we imposed affect not only mitochondria, but all of the other organelles and cellular components with which they interact. A simple way this could be done systematically would be to reclone the gTEMP reporters (or better, the improved B-gTEMP version) for targeting to other compartments, followed by careful validation of targeting (e.g. to nucleus, ER lumen, mitochondria-associated membranes (MAMs), peroxisomes, endosomes, Golgi, the plasma membrane and cytosol. Such a study would provide a picture of how alterations in mitochondrial metabolism and heat production affect intracellular temperature and its regulation more globally. An alternative approach would be to employ the recently developed palette of small-molecule intracellular temperature reporters, the Thermo Greens^35^. Earlier studies using temperature-sensitive dyes targeted specifically to the ER^36–38^ have indicated how the temperature of this organelle responds to external stresses and physiological stimuli. Use of the B-gTEMP reporters targeted to different submitochondrial compartments would also shed light on heat transfer within mitochondria, and how this varies with organelle architecture in different cell types. The use of reporters in isolated mitochondria might also provide insight into how much buffering of heat transfer is provided by the cytosol, although this would not address the temperature of the organelle itself *in vivo*. Note, however, that not all such dyes and reporters may be employed simultaneously, although studies in parallel should help address many of the outstanding questions. Since every method has technical limitations, it is obviously desirable that all conclusions arising from our and similar studies should, as far as is feasible, be independently verified using different approaches.

We previously reported^6^ that isolated OXPHOS complexes I and V are heat labile, though OXPHOS complex II (cII), cIII and cIV activities have temperature optima in the 50-55 °C range when studied in intact mitochondrial membranes. Others have recently reported that the imposition of an external temperature of 43 °C or above to cells or to isolated mitochondria damages respiratory function and disturbs OXPHOS complex and supercomplex organization^32^. Thus, externally applied heat delivered to mitochondria, which our assays indicate are already at elevated temperatures, may push them beyond their operational limit, implying that mitochondrial heat production *in vivo* must be finely attuned to cellular needs, to avoid catastrophic overheating. On the other hand, a recent meltome analysis of human proteins found that those constituting the respiratory chain complexes have median Tm values of around 60 °C (Ref. 39), consistent with our inference of high mitochondrial temperature.

### Inhibition of each OXPHOS complex produces specific temperature effects on mitochondria

As the OXPHOS system is traversed from cI through cV, we observed significant increases in the temperature effect of specific inhibitors, with essentially the same pattern observed in two iMEF lines, U2OS, HEK293T and even in *Drosophila* S2 cells, as far as methods allowed, i.e., oligomycin> KCN > antimycin ≥ rotenone. Oligomycin inhibits all protonmotive components of the OXPHOS system, including those that feed electrons via ubiquinol to cIII, even though they are not directly protonmotive. The lesser amount of mitochondrial temperature decrease produced by blocks on cI, cIII or cIV (Fig. 2) is not the result of incomplete inhibition: the inhibitor doses were pre-checked on all cell-lines by respirometry and are similar to those used in many other studies.

Instead, we posit that it reflects the proportion of electron flow entering the respiratory chain at each step. Since rotenone and antimycin inhibition had the same mitochondrial temperature effect in iMEFs, both individually and in combination, we infer that electron flow entering the chain independently of cI in this cell background is negligible. In contrast, antimycin produced a significantly greater mitochondrial temperature decrease in U2OS cells than did rotenone (Fig. 2B), from which we conclude that around 20-30% of electron flow into cIII in this cell line must come from other ubiquinone-linked enzymes. In iMEFs (Fig. 2C, 2D) and S2 cells (Fig. 2E), though not in HEK293T or U2OS cells, KCN had a significantly greater mitochondrial temperature effect than antimycin, which implies that, in these cell lines, a measurable amount of electron flow to oxygen enters the OXPHOS system via cytochrome *c* independently of cIII, i.e. via the Mia40-Evr1/ALR protein import pathway, albeit that we were not able to verify this directly using the Evr1/ALR inhibitor MitoBloCK-6. There may also be other, as yet uncharacterized, electron donors to cytochrome *c*. In all cell lines that we were able to test, oligomycin produced a significantly greater mitochondrial temperature decrease than KCN (Fig. 2A-2D). Since KCN should block all respiratory proton pumping across the IMM, we infer that cV itself must be able to generate some heat in the complete absence of cellular respiration, presumably by the hydrolysis of mitochondrially imported ATP. Since this residual thermogenesis is inhibited by oligomycin, it must nevertheless involve cV, in a reaction that is itself protonmotive, and hence limited by the rate of dissipation of the IMM proton gradient by other processes, such as protein, metabolite and ion transport.

In cells expressing AOX, the initial effects of OXPHOS inhibition at these various steps was different (Fig. 6A, 6B). Rotenone inhibition of electron flow through cI initially produced the same temperature decrease in AOX-expressing iMEF mitochondria as in control iMEF mitochondria (Fig. 6A, 6B). This is consistent with the fact that AOX by-passes cIII+cIV but not cI^22, 40^. In contrast, the mitochondrial temperature decrease produced by antimycin inhibition of cIII was substantially less than in control cells (Fig. 6B). Electron flow through the inherently thermogenic AOX is expected substantially to replaces the normal heating effect of electron flow through cIII+cIV. Note that, when cIII and cIV are able to function normally, electrons flowing through cI should not be diverted via AOX^23^. However, the mitochondrial temperature decrease brought about by antimycin in AOX-expressing MEFs was not zero. Part of the explanation is that the capacity of AOX to support electron flow from cI through to oxygen in these cells remains below that of cIII+cIV, as inferred previously by respirometry^26^. Furthermore, the full engagement of AOX for cI-linked substrates^23^ requires the accumulation of ubiquinol, the reduced form of coenzyme Q. Nevertheless, the initial temperature decrease produced by antimycin in the mitochondria of AOX-expressing cells is maintained only for a relatively short time, approximately 10 min (Fig. 6C), before rewarming commences. The fact that the initial effect of KCN in AOX-expressing cells was greater than that caused by antimycin supports the idea that some of the electron flow through cIV in iMEFs is introduced via cytochrome *c* independently of cIII, for which AOX does not provide a by-pass.

### High mitochondrial temperature is maintained under diverse metabolic stress conditions

Despite the fact that. under the rather extreme conditions of toxic inhibition of the OXPHOS complexes, high mitochondrial temperature is compromised, we uncovered several contexts wherein it appeared to be homeostatically maintained, hinting at one or more metabolic adaptation processes. First, it appears to decline only slowly and minimally in response to nutrient starvation, and to be restored by renewed substrate availability (Fig. 5), consistent with previous estimates by an independent method, using so-called Upconversion nanoparticles^41^. This implies the existence of a metabolic reserve even in cells such as fibroblasts that are not normally considered repositories for storage molecules such as triglycerides or glycogen. Second, in cells expressing AOX we recorded a gradual restoration of high mitochondrial temperature after the initial decrease provoked by inhibition of the OXPHOS complexes (Fig. 6A, 6C). Although AOX is not normally present in vertebrate cells, the metabolic flexibility that it confers may be important in maintaining mitochondrial temperature and organismal viability in the vast array of species that do possess AOX but are subject to wide external temperature fluctuations, namely bacteria, fungi, plants, eukaryotic protists and many animals as well^40^. The exact nature of the metabolic changes that enable AOX-endowed cells to recover normal mitochondrial temperature over periods of 40-70 min remains to be elucidated and may be complex. Logically it should involve a remodeling of metabolism towards the oxidation of flavin-linked substrates that directly reduce ubiquinone.

Rewarming was evident in the case of cells in which cI was inhibited by rotenone (Fig. 6A, 6C), indicting that it does not involve the acquisition of additional capacity for AOX to receive electrons via cI. Rather, it must involve a metabolic switch to other pathways, even though via AOX these do not support ATP production, only heat generation.

The rather minimal effect of BKA, i.e. a possible transient, small increase in mitochondrial temperature, followed by an equally modest decrease, was also surprising. BKA treatment of immortalized fibroblasts has previously been reported to induce a switch to glycolytic ATP production and decreased flux through the respiratory chain, whilst maintaining high mitochondrial membrane potential^21^. The modest mitochondrial temperature decrease that was inferred may simply correspond with that seen in cells deprived of substrate over the time-period of observation (compare Fig. 5A and 5E or 5B and 5F). Thus, the efficient exchange of adenine nucleotides between mitochondria and the cytosol is not required for the maintenance of high mitochondrial temperature. Instead, the usually minor contribution of intramitochondrial ATP turnover can be boosted sufficiently to drive mitochondrial heat production to near normal levels, possibly involving, once again, a contribution from cV-linked ATP hydrolysis.

The most surprising result that we obtained was the maintenance of high mitochondrial temperature in cells subjected to prolonged treatment with anisomycin at a dose in excess of that which profoundly inhibits cytosolic protein synthesis^42, 43^. Protein synthesis uses between 35-70% of the ATP produced in cells^44–47^. We therefore expected that the predicted decrease in ATP demand would lead to decreased flux through the respiratory chain and to a sustained drop in mitochondrial temperature. Instead, anisomycin treatment led to a modest but transient increase (∼4°C) in mitochondrial temperature (Fig. 7A), with gradual reversion to the original mitochondrial temperature after 2-3 h in the presence of the drug. Even after prolonged (18 h) anisomycin treatment, leading to widespread cell death, high mitochondrial temperature was maintained in the surviving cells. The same effects, but without cell death, were also seen in AOX-expressing cells (Fig. 7B).

Similarly, overnight growth in media that are predicted to stimulate mitochondrial as opposed to cytosolic ATP production had no effect on mitochondrial temperature (Fig. S3B). Once again, the precise metabolic changes that underlie these phenomena remain uncertain and may not be easy to dissect. Nevertheless, our observations imply a homeostatic mechanism adjusting mitochondrial heat production to metabolic supply and demand, but keeping mitochondrial temperature within a narrow range.

Based on the inhibitor studies shown in Fig. 7C and 7D, mitochondrial temperature maintenance in anisomycin-exposed iMEF cells is less and less dependent on cI with time, yet remains fully dependent on cV. This suggests that, as cellular ATP usage dwindles, cV may start to operate thermogenically as an ATPase, although this does not explain how the resulting proton gradient across the IMM is dissipated. A more complex mechanism, such as the futile creatine cycle recently documented in brown fat^48^, may need to be invoked. The adenine nucleotide carrier (ATP-Mg/Pi), encoded by a small multigene family, which ensures the net delivery of adenine nucleotides to the mitochondrial matrix under specific physiological conditions^49^, may also be involved in sustaining a futile cycle of intramitochondrial ATP hydrolysis, as well as possible reverse electron transport through cI. In anisomycin-exposed, AOX-expressing cells, where AOX itself provides an alternative mechanism of thermogenesis, cells remained more dependent on cI for the maintenance of high mitochondrial temperature over time, but less dependent on cV.

An important caveat of our findings is that we cannot formally distinguish whether metabolic adjustments are being made in response to external stresses, with mitochondrial temperature maintenance just a ‘by-product’ of these changes or, conversely, that mitochondrial temperature is directly responsive to external stresses, entraining metabolic adjustments in its wake. A search for an endogenous mitochondrial temperature sensor and an associated signal transduction machinery may help resolve this.

### Theoretical considerations

A mitochondrial temperature in excess of 50 °C places severe constraints on the mechanisms of intracellular heat conductance. Here we postulate that, at least in mammals, mitochondrial heat production sustains the cell broadly at around 37 °C, but must be responsive both to external heat stresses and to metabolic changes taking place within the cell, including those within the mitochondrion itself. Thus, the molecular and physical mechanisms that govern heat flow within and out of mitochondria must be both robust and flexible.

As Fahimi and Matta have pointed out^50^, the thermal conductivity of the mitochondrial membranes must be very small at the distances encountered in the cell between the hot and cold compartments. Fahimi and Matta^50^ termed this the mitochondrial paradox and offered a physical explanation based on spikes in local temperature associated with proton passage from the IMS to the matrix. They posit that the dehydration and deprotonation of the hydronium ion on the IMS side and the reverse process on the matrix side creates a mode of heat transfer consistent with the time scale of these processes and with the very low thermal conductivity of ATP synthase. Based on these considerations they were able to predict a 7K difference across the membrane. To our knowledge, their work is the only physically sound hypothesis yet proposed, to account for the surprising temperature differences inferred experimentally, and whose validity has been widely debated^13, 31, 51–53^.

Matta and colleagues have also argued, based on thermodynamic principles^54^, that the true efficiency of ATP synthase must be close to 90%, taking account of heat dissipation by informational transactions (Landauer’s principle^55^) which is compensated by the inherent electrostatic potential of the enzyme^56^. This may partly explain why the additional temperature decrease produced by inhibiting ATP synthase in addition to inhibiting respiration is both small, but non-negligible.

### Biological implications of high mitochondrial temperature

Although the amount of mitochondrial temperature decrease produced after OXPHOS inhibition by oligomycin, when measured by the MTY and mito-gTEMP methods, was similar, it was not identical, and the rather small difference was judged to be significant for at least two of the cell-lines studied, U2OS osteosarcoma cells and iMEFs. Although both probes are efficiently targeted to mitochondria, they appear to be localized in different submitochondrial compartments. As derivatives of GFP, both of the mito-gTEMP reporter polypeptides linked to the COX8 targeting peptide should reside in the mitochondrial matrix^18, 57^. Conversely, MTY is a rhodamine-related lipophilic dye whose uptake requires a mitochondrial membrane potential, and which is rapidly lost from mitochondria when the membrane potential is abrogated^6^. Its localization at or very close to the inner face of the IMM is supported by our sub-mitochondrial fractionation experiments, showing that it co-fractionates preferentially with SMPs rather than with the mitochondrial matrix-enriched fraction. This, despite reports that at least some of it interacts with the matrix protein ALDH2 (Ref. 19). Since the IMM is also the location of mitochondrial heat generation, it should logically be slightly warmer than the matrix that it surrounds, accounting for the small temperature differential implied by the two methods.

Since high mitochondrial temperature was here documented in five different cell-lines from three species of animal, including a poikilotherm, it is clearly not just an exceptional feature of one cell-type. Rather it would appear to be a common property of mitochondria and of the chemistry of the OXPHOS system. Moreover, it suggests a novel role for the universal double membrane system of mitochondria, as an insulating layer enabling mitochondria to maintain a high internal temperature, whilst allowing the remainder of the cell to operate at a lower temperature. If this is correct, a corollary is that the ratio of mitochondrial volume to surface area may change in response to metabolic needs and physiological signals that regulate mitochondrial heat output. Put more simply, mitochondria might be expected to fragment into smaller entities when heat output rises, so as to maintain internal mitochondrial temperature, and to fuse into filamentous structures when heat output is low. They may also fragment if mitochondrial heat production is simply abolished and mitochondria need to absorb as much heat as possible from the rest of the cell, in order to maintain a minimal metabolism.

This proposition could account for some of the apparently paradoxical dynamics of mitochondria which, when subject to various stresses can either undergo widespread fission or hyperfusion^58^. The mitochondrial fusion/fission cycle has hitherto been associated primarily with mitochondrial quality control^59^, which is disturbed in patholological states and during aging^60^. However, mitochondrial temperature homeostasis could also be a determinant. Accordingly, it has recently been reported that inhibition of cV, but not of the other OXPHOS complexes, triggers mitochondrial fragmentation^61^. Treatment with rotenone or antimycin were reported to result in rapid depletion of intramitochondrial ATP^61^, consistent with our inference that these inhibitors allow some residual mitochondrial heat output driven by ATP hydrolysis, whilst oligomycin does not. Using mito-gTEMP, it was previously shown that treatment of cells with an uncoupler, which is known to provoke mitochondrial fragmentation^62^, also resulted in a mitochondrial temperature increase of ∼6 °C (see Fig. 4 of Ref. 7). Using a different spectrophotometric method, other authors estimated the effect of a chemical uncoupler in the 4-5 °C range^34^. Consistent with the idea that mitochondrial fusion is associated with low mitochondrial heat output, the cell line with the greatest inferred temperature difference between the two methods used in the present study to assess mitochondrial temperature, and also the one where this difference was the most variable experimentally, was U2OS (Fig. 3E), a cell line in which mitochondria are mostly filamentous, e.g. see Ref. 63. In other cell-lines, such as iMEFs, mitochondria are highly fragmented under standard culture conditions, e.g. see Ref. 64.

Staining of cells with MTY revealed bright puncta within mitochondria (Fig. 4B), the nature of which is unknown. One possibility is that these may correspond with nucleoids. Their bright staining could indicate that they are cooler than their mitochondrial surroundings, or could be an artefact, due to binding of the dye to one or more components of the nucleoid. In *Saccharomyces cerevisiae* the orthologue of ALDH2 (Ald4) is a nucleoid protein, but this has not been reported to be the case in mammals^65^.

Since the OXPHOS system is of bacterial origin, its heat output in bacteria should be similar to that of mitochondria, and we would expect that the thermal insulation provided in gram-negatives by the double membrane and lipopolysaccharide (LPS) cell wall, or by the thicker cell wall in gram-positives, should be at least as effective as that of the double membrane of mitochondria. It will therefore be interesting to implement thermosensitive reporters such as gTEMP or B-gTEMP in bacteria, to assess intracellular temperature, which we would predict to be once again some 15-20 °C warmer than ambient temperature. This has important potential implications for understanding the metabolic biochemistry of bacteria, including antibiotic action.

Similarly, considering that heat generation is an inescapable cellular function of metabolically active mitochondria, a loss of this activity, as in cases of mitochondrial disease, may compromise the maintenance of intracellular temperature or induce futile cycles of cytosolic ATP generation and usage that may have profound metabolic effects. This could be a crucial aspect of mitochondrial pathology^66^ and may also influence progression in other metabolic diseases, including cancer^67^. Some of these and related issues are discussed in the recent paper of Fahimi and colleagues^68^.

## Conclusion

On the basis of our findings, we propose that the maintenance of high mitochondrial temperature should be regarded as an important homeostatic process of the organelle, along with more traditionally considered parameters such as membrane potential, pH, ATP, metabolite levels and ROS.

## MATERIALS AND METHODS

### Cell culture

All cell-lines used are freely available and are not subject to any ethical restrictions on use. Human embryonic kidney-derived cells expressing a mutant version of the SV40 large T antigen, HEK293T (Ref. 69), U2OS (2T) osteosarcoma cells^70^, two isolates of immortalized mouse embryonic fibroblasts, iMEF(L7)^26^ and iMEF(P)^25^ and immortalized mouse embryonic fibroblasts expressing *C. intestinalis* alternative oxidase, iMEF(AOX)^25^ were cultured in DMEM medium containing 4.5 g/L glucose, 10% heat inactivated FBS, and 100 U/mL each penicillin and streptomycin, at 37 °C in 5% CO_2_ with 95% humidity. Cells were also grown in low-glucose medium (Sigma, D5546) containing 1 g/L glucose, supplemented with 25 mM galactose where indicated. *Drosophila melanogaster* S2 (Schneider 2) cells (Invitrogen), originally derived from a primary culture of late stage (20-24 h old) embryos^71^, were cultured in suspension in Schneider’s medium (Sigma S9895-1L) supplemented with 10% heat-inactivated FBS, 100 U/mL each penicillin and streptomycin at 25 °C and diluted 1:6 every 3-4 days.

### Transfection of mammalian cells with mito-gTEMP

To avoid the use of neomycin/geneticin selection, HEK293T cells already being resistant, the mito-gTEMP ratiometric reporter pair (see Fig. 3A) was recloned from mito-gTEMP_pcDNA3 (Addgene, plasmid #109117) into the pTriEx™-1.1 Hygro plasmid (Novagen 70928). Stably expressing cell lines were generated using FuGENE® HD Transfection Reagent (Promega), according to manufacturer’s instructions. Based on preliminary trials, hygromycin selection was implemented at the following doses: iMEF(P) and U2OS – 145 µg/mL, iMEF(AOX) 240 µg/mL, and HEK293T 450 µg/mL.

### Spectrofluorometry

For spectrofluorometric measurement of mitochondrial temperature using Mito Thermo Yellow, MTY^19^, sub-confluent cells were treated with 100 nM MTY supplemented with warmed fresh medium, for 15 min. Cells (5 x 10^6^ HEK293T and U2OS cells, 7 x 10^6^ iMEFs, and 10 x10^6^ S2 cells) were trypsinized and collected by centrifugation at 250 g_max_ for 3 min. The pellet was washed once in 10 mL PBS warmed to 37 °C, then maintained as a concentrated pellet at 37 °C for 10 min, for anaerobiosis. Cells were resuspended in PBS in a magnetically stirred 3.5 mL quartz cuvette with 10 mm optical path (Hellma Analytics, Germany), placed in a Peltier temperature-controlled chamber set at 38 °C (25 °C for S2 cells). The time-based fluorescence signal at constant emission (excitation 542 nm, emission 562 nm) was generated using a QuantaMaster^TM^-6/2003 LPS-220B spectrofluorometer (PTI-Horiba, Japan, Fig. S1). For ratiometric spectrofluorometry of cell lines genetically expressing mito-gTEMP, cell preparation and collection were carried out similarly, with the emission of both fluorophores and their ratio recorded over time, with excitation at 360 nm and emission for Sirius at 425 nm and for mT-Sapphire at 509 nm.

### Ratiometric temperature calibration for mito-gTEMP *in vivo*

Cells were prepared as described above and, once resuspended in the cuvette, were treated with 5 μM oligomycin to block ATP synthesis, hydrolysis and respiratory electron transfer. As soon as the fluorescence ratio reached a steady-state, the temperature of the Peltier jacket holding the cuvette was shifted to 30 °C and then in 3 °C steps to 42 °C, in each case recording the fluorescence ratio during the third minute. The experiment was repeated (n=9) and the ratios of mT-Sapphire/Sirius fluorescence were plotted against temperature to generate a linear calibration function.

### Sub-mitochondrial fractionation

iMEFs were labelled with MTY as above, on 2 x 20 cm plates, with unlabelled cells treated in parallel. Following trypsinization, cells were washed with 10 mL of ice-cold PBS and mitochondria isolated by homogenization under hypotonic conditions, as described^72^. Mitochondria were resuspended in 2 mL of hypotonic buffer (5 mM HEPES, 2 mM EDTA, 1 mM PMSF, pH 7.2) plus 5 mM each of pyruvate, glutamate, and malate (PGM) mix. Following incubation for 5 min on ice, digitonin was added to 1% (v/v), with gentle shaking on ice for 15 min, followed by centrifugation at 20,000 g_max_ for 10 min at 4 °C (Centrifuge 5810 R, Eppendorf)^73–75^ to separate mitoplasts (pellet) from outer mitochondrial membrane and inter-membrane space (IMS) components (supernatant).

For estimating fluorescence, the pellet was resuspended in the hypotonic buffer plus PGM to the same volume (2 ml) as the supernatant, and both samples were brought to 38 °C before fluorescence measurement (excitation 542 nm, emission 562 nm), which was carried out over 1 min for all samples to verify that it had reached a constant value. In a further fractionation step, mitoplasts were sonicated on ice at high power, using five 30 s on/off cycles, i.e. for 5 min (Bioruptor Sonicator, Diagenode)^73–75^, then centrifuged at 27,000 g_max_ for 15 min at 4°C (Optima XPN-100 Ultracentrifuge, Beckman Coulter, with SW60 Ti rotor). Resuspension of the pellet (sub-mitochondrial particles, SMP) in the same volume as the supernatant (matrix fraction), temperature equilibration at 38 °C and fluorescence measurements were performed as above. The autofluorescence of control cell samples was subtracted from the fluorescence of the corresponding fraction of MTY-labelled cells, enabling the proportionate recovery of label in different fractions to be estimated. See Fig. S2H for a summary of the fractionation scheme.

### Respirometry

In order to determine the exact concentrations of inhibitors required for complete inhibition of each OXPHOS complex, we used the same amount of cells as used for spectrofluorometry to measure oxygen consumption with a Clark-type electrode (Oxytherm system, Hansatech, UK). Intact cell respiration was recorded from cells suspended in 500 µL of Respiraton Buffer A (225 mM Sucrose, 75 mM mannitol, 10 mM Tris/HCl, 10 mM KCl, 10 mM KH_2_PO_4_, 5 mM MgCl_2_, pH 7.4) plus 10 mg/mL freshly added BSA, permeabilized by the addition of 80 µg/mL digitonin at 37 °C. Substrate concentrations were as follows; 10 mM ADP, 5 mM pyruvate and malate (cI+cIII), 10 mM succinate (cII+cIII), 50 mM TMPD/1 mM ascorbate (cIV). The required concentrations for full inhibition were determined to be 3 µM rotenone (cI inhibitor), 1 µM antimycin (cIII inhibitor), 0.8 mM potassium cyanide (KCN, cIV inhibitor), 5 µM oligomycin (cV inhibitor), and 100 µM n-propyl gallate (AOX inhibitor).

### Thermo-profiling of mitochondria in cells treated with OXPHOS inhibitors

Cells were prepared for spectrofluorometry, and inhibitors were added to the above concentrations, once fluorescence or fluorescence ratio had reached a steady state, generally 10 min after initial oxygenation. Fluorescence values were then studied over time, until a new equilibrium value was reached, but only until 35 min for mito-gTEMP. MTY fluorescence was calibrated against temperature in each experiment, by raising and lowering the Peltier by 3 °C at the end of the experiment, and recording the fluorescence values reached. The change in temperature was then estimated using this internal calibration. For estimating temperature changes using mito-gTEMP, the calibration function described above was applied to the actual values obtained for the mT-Sapphire/Sirius ratio. The adenine nucleotide transporter (ANT) inhibitor Bongkrekic acid (Sigma, B-6179) and ALR/Evr1 inhibitor MitoBloCK-6 (Calbiochem, Sigma-5.05759) were used similarly at 100 µM.

### Cytoplasmic protein synthesis inhibition

Cells were seeded and, after 24 h, the medium was replaced with otherwise complete medium without penicillin or streptomycin, but with 150 µM anisomycin (A-9789, Sigma). Cells were incubated for various times up to 18 h. For measuring the mito-gTEMP fluorescence ratio, cells were then treated as for control (untreated) cells expressing mito-gTEMP, except that for all time-points except that at t=0, the cells were resuspended in PBS containing 150 µM anisomycin. For measuring MTY fluorescence in anisomycin-treated cells, 100 nM MTY was added 15 min prior to each time point, followed by resuspension in PBS containing 150 µM anisomycin and the addition of OXPHOS inhibitors as for control (untreated cells).

### Microscopy

Cells were seeded on glass bottom dishes (35 mm MatTek, USA) and grown for 24 h in standard growth media at 37 °C, 5% CO_2_. The culture medium was replaced with prewarmed medium containing fluorescent dyes, namely 100 nM MTY and/or 100 nM MitoTracker™ Deep Red FM (ThermoFisher Scientific). After 15 min incubation, the staining medium was removed, and cells washed once with fresh, prewarmed PBS then kept in PBS at 37 °C until analyses were complete. The expression and localization of mT-Sapphire in transgenic cell lines was imaged by spinning disc (X-light V2, CrestOptics) confocal fluorescence microscopy (Nikon FN1), with excitation at 360 nm (Cool-Led pE-400 light source) and emission filter 525/50 nm, with Nikon 60X/1.0 dip objective (CFI Apochromat NIR 60X W). Laser excitation light intensity was adjusted to minimize photobleaching. Fluorescence images for MitoTracker^TM^ Deep Red FM (excitation 633 nm, emission 700/75 nm) were generated similarly. Live-cell and time-lapse images of cells stained with MitoTracker^TM^ Deep Red FM or MTY (excitation 542 nm, emission 562 nm) were captured by laser-scanning microscopy (Nikon A1R), using a Nikon 60x/1.27 water-immersion (CFI SR Plan Apo IR 60XC WI) objective. The detector sensitivity was adjusted for each sample to optimize the image brightness and to avoid saturation. The Nikon A1R with N-SIM laser scanning confocal microscopy system includes a live-cell supporting chamber where CO_2_, temperature of the culture dish (surrounded by heated-water supply), and temperature of objective in use can be controlled.

Time-lapse imaging was performed on cells maintained at 37 °C with one image /10sec for 10 mins, with a 2 min pause for addition of oligomycin to 5 µM, followed by a further 30 min of image capture. Signal analyses of time-lapse images were performed with ImageJ (Fuji), by surface-plot analysis.

### Statistics

All data are presented as means ± SD. Statistical significance was calculated by standard unpaired one-way ANOVA with Tukey HSD test (www.astatsa.com) or, where appropriate, by Student’s t test (LibreOffice Calc). Values of p < 0.05 were considered significant.

### Image presentation

Micrographic images and fluorescence traces were cropped, rescaled and optimized for brightness and contrast, with labels added or removed, but with no other modifications.

## ACKNOWLEDGMENTS

We thank Malgorzata Rak and Eric Dufour for useful discussions, Kateryna Gaertner for assistance with subcellular fractionation, Tea Tuomela for technical assistance, Maria Carretero-Junquera for artwork and the Biocenter Finland-supported Tampere Imaging Facility for access to its infrastructure.

## AUTHOR CONTRIBUTIONS

M. Terzioglu = Conceptualization; Methodology; Validation; Formal analysis; Investigation; Resources; Data curation; Writing – original draft preparation; Writing – review & editing; Visualisation; Supervision

K. Veeroja = Conceptualization; Methodology; Validation; Investigation; Writing – review & editing

T. Montonen = Methodology; Investigation; Resources; Writing – review & editing; Visualisation Teemu O. Ihalainen = Conceptualization; Methodology; Resources; Writing – review & editing; Visualisation; Supervision

T. S. Salminen = Conceptualization; Resources; Writing – review & editing; Supervision; Project administration; Funding acquisition

P. Bénit = Conceptualization; Methodology; Writing – review & editing

P. Rustin = Conceptualization; Methodology; Writing – review & editing

Y.-T. Chang = Conceptualization; Methodology; Resources; Writing – review & editing; Project administration; Funding acquisition

T. Nagai = Conceptualization; Methodology; Resources; Writing – review & editing; Project administration; Funding acquisition

H. T. Jacobs = Conceptualization; Methodology; Validation; Formal analysis; Resources; Writing – original draft preparation; Visualisation; Supervision; Project administration; Funding acquisition

## LIST OF SUPPLEMENTARY FILES

temp SI.pdf – 7 supplementary figures and their legends

Suppl movie S1.avi – supplementary movie

## SUPPLEMENTARY INFORMATION

## LEGENDS TO SUPPLEMENTARY FIGURES

**Supplementary Figure S1.**
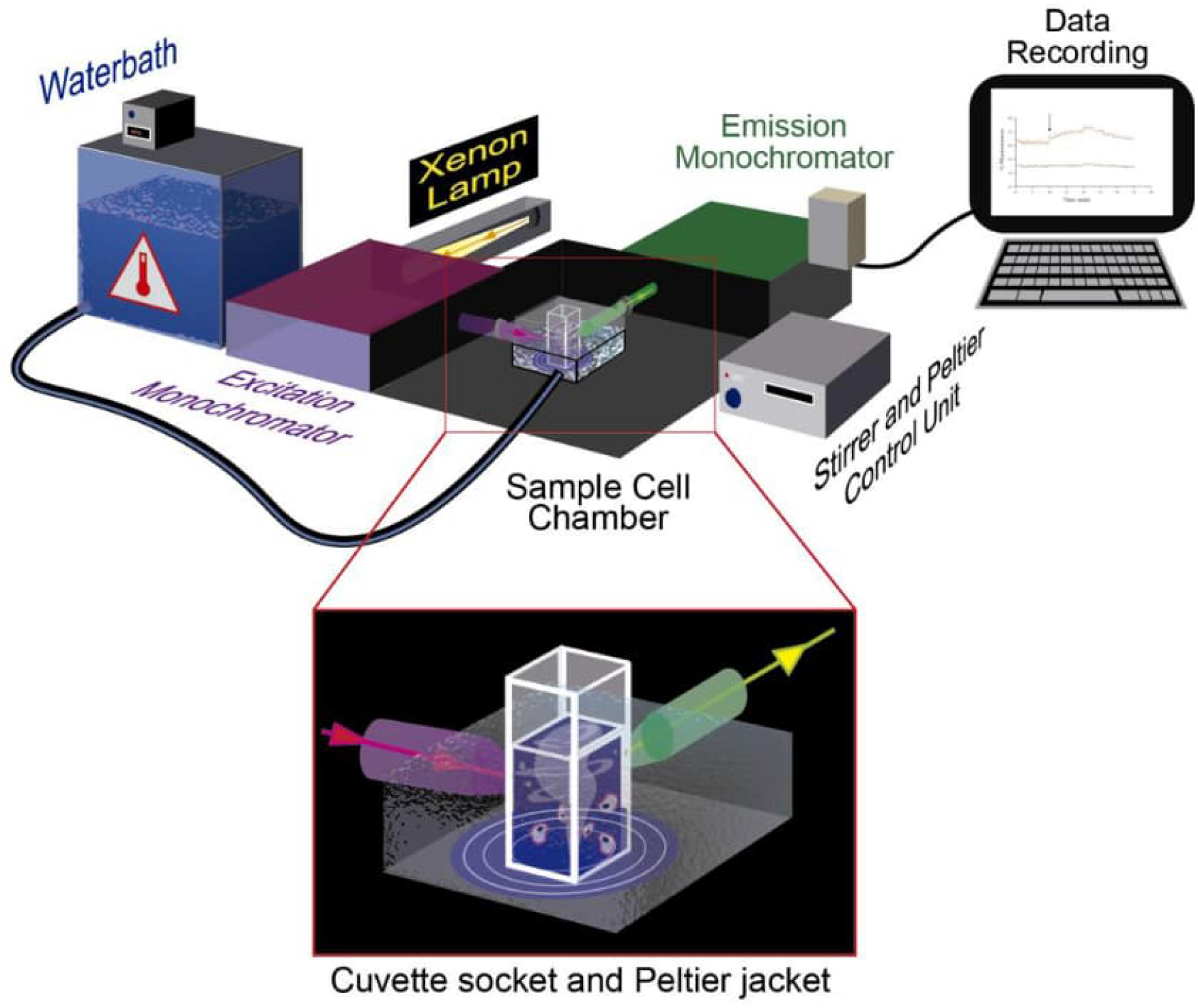
Instrumentation for mitochondrial temperature estimation using fluorescent probes. Schematic illustration of spectrophotometer, QuantaMaster^TM^ QM-6/2003 LPS-220B with a Peltier temperature-controlled sample cell chamber. Waterbath (Julabo® CORIO^TM^ CD, Germany) with refrigerated/heated circulator (-20 to 150 °C) provides accurate temperature range via circulation around the sample cell chamber. The xenon lamp (Xenon short arc UXL 75XE, USHIO, Japan) used as the light excitation source. The excitation monochromator selects the desired wavelength of excitation light, which is focused at the sample position. The emission monochromator is scanned across the desired emission range and fluorescence intensity is recorded in the detector as a function of emission wavelength. While data is recorded, the cell sample is stirred continuously, maintaining oxygen concentration far above the level required for fully active cellular respiration. For instrument validation see Fig. S2. Illustration by Maria Carretero-Junquera.

**Supplementary Figure S2.**
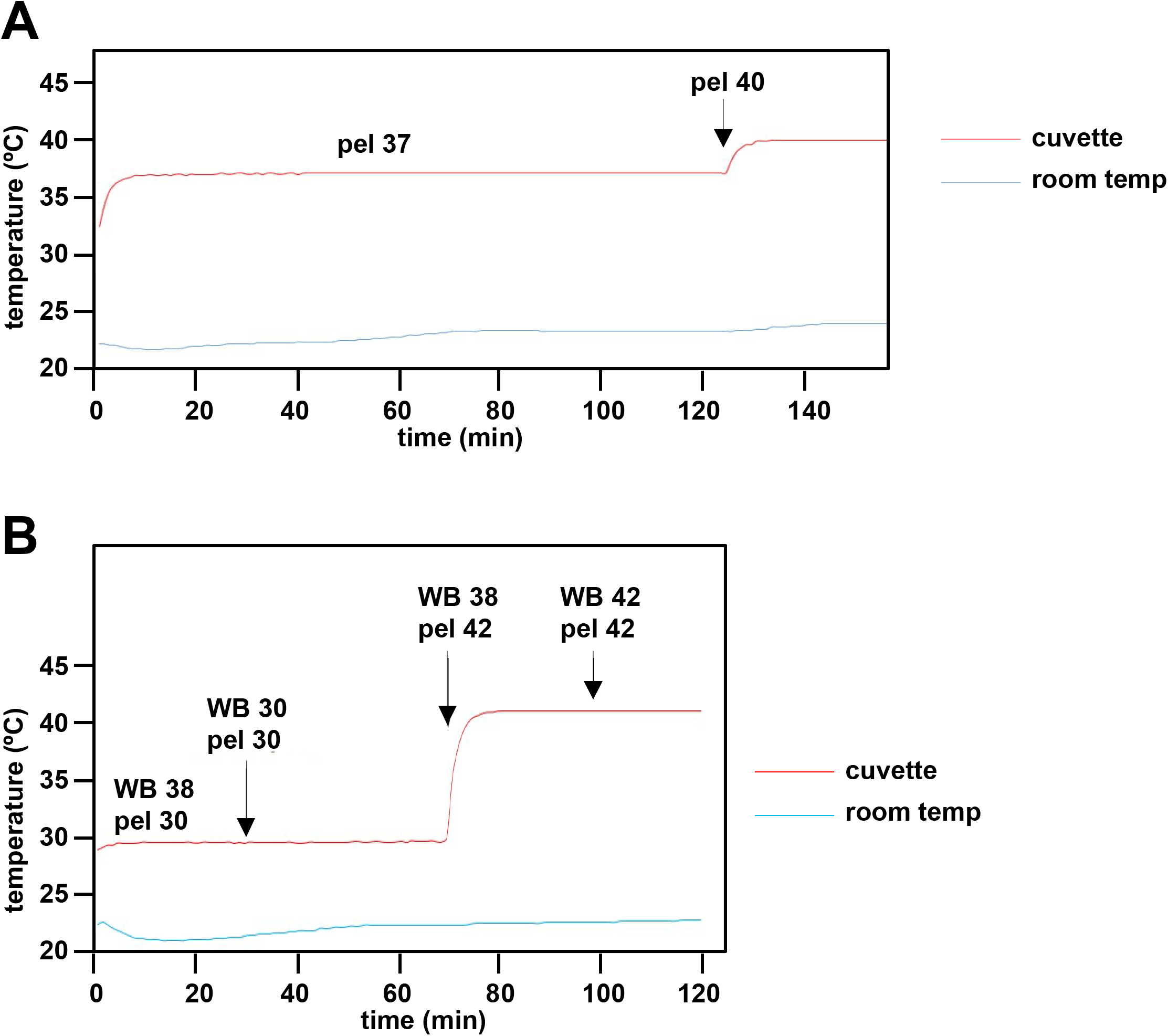
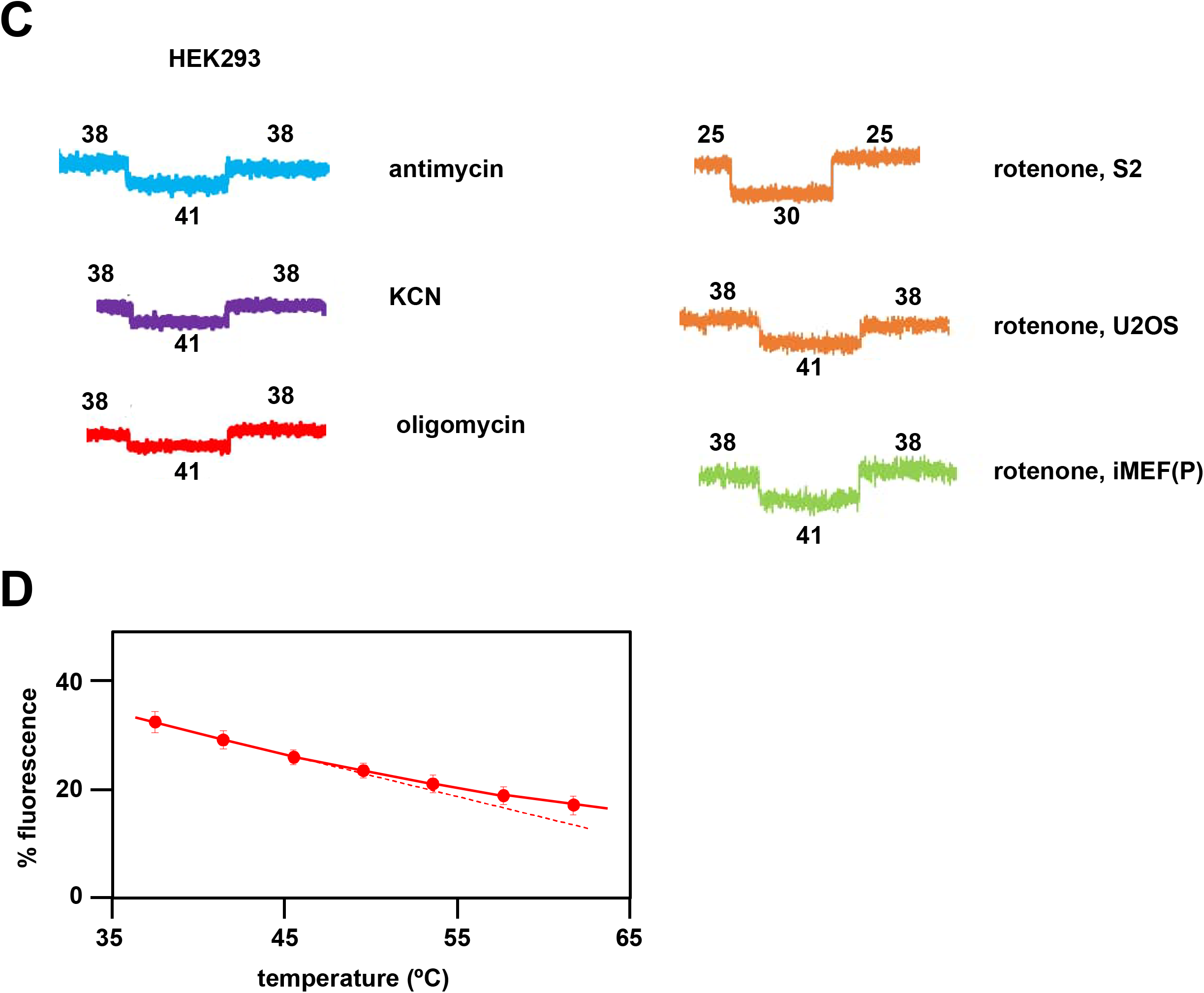
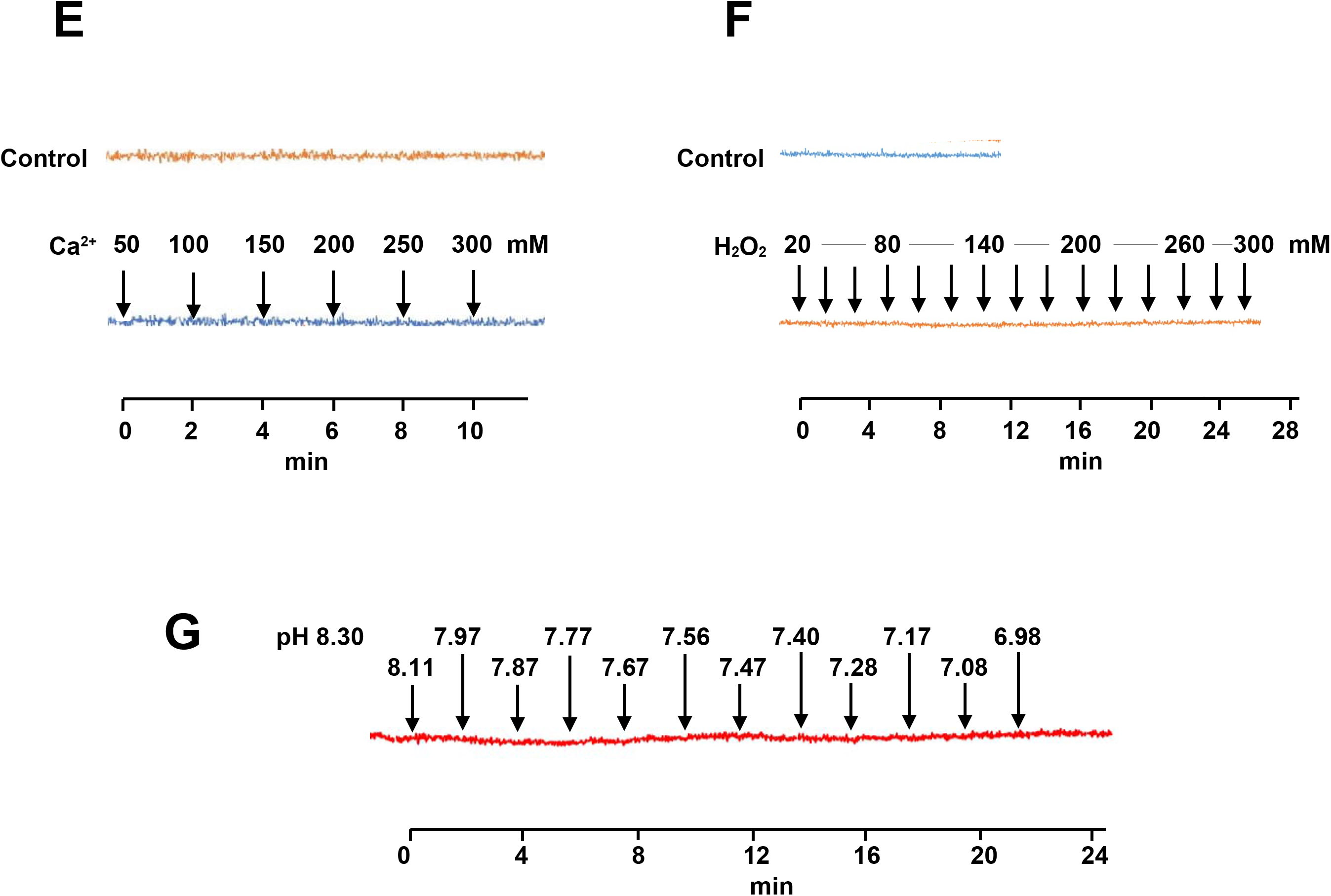
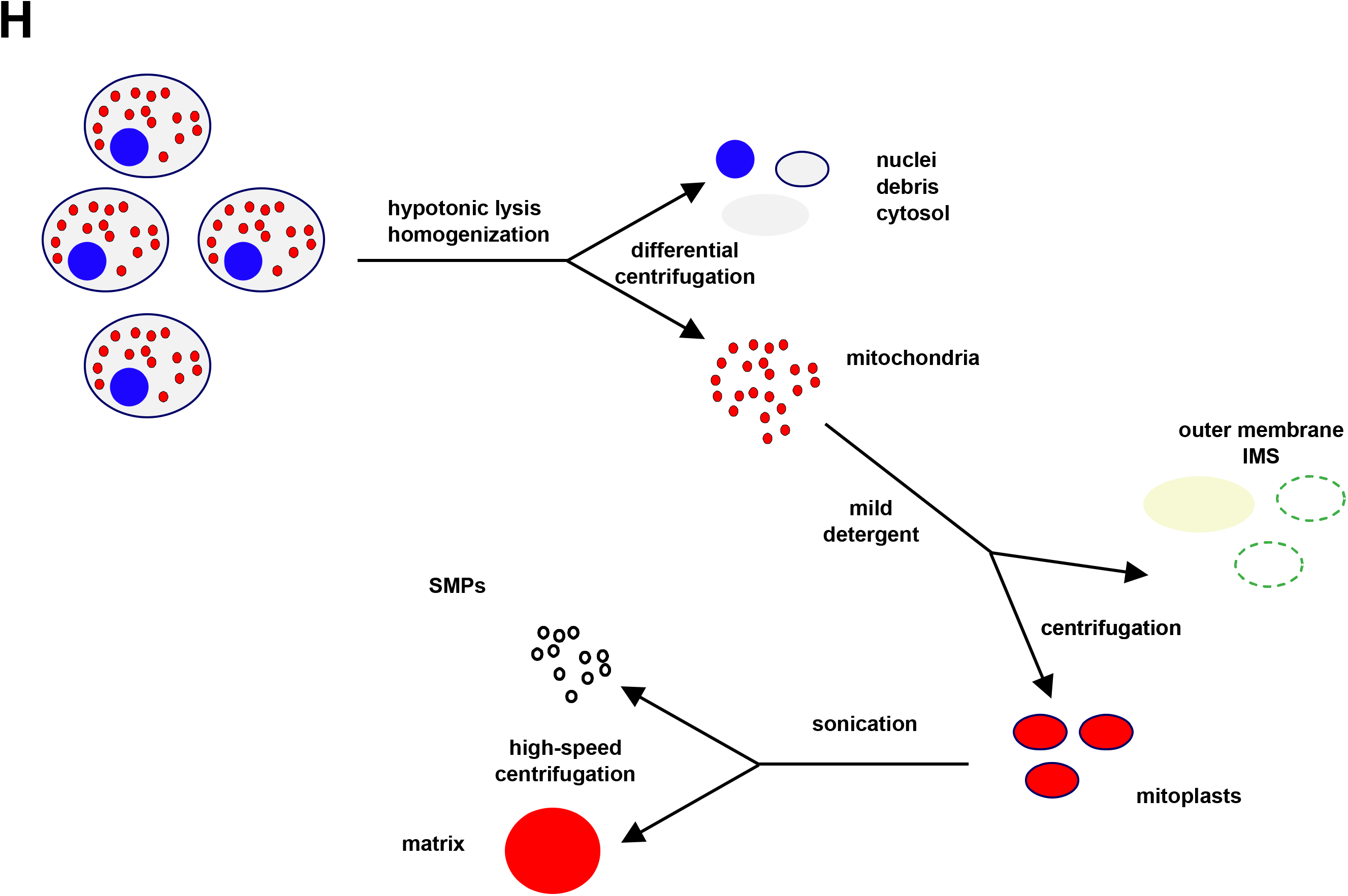
Instrument and fluorescence validation steps for estimation of mitochondrial temperature. (A) In an initial step, an external temperature probe was used to validate the temperature readings of components of the apparatus (water-bath, Peltier, cuvette) as shown on the digital displays. After a 3 °C temperature shift, the Peltier took ∼2 min to stabilize at the new temperature, and held steady despite minor fluctuations in the external room temperature. (B) The cuvette temperature was not affected by water-bath temperature changes or differences from the set temperature of the Peltier. The water bath temperature (not shown) itself took ∼5-6 min to restabilize. Note that the cuvette temperature in steady-state was 1 °C lower than that of the Peltier, which was taken account of in the calibration shown in Fig. 3D. (C) Sample traces showing calibration steps after fluorescence of the indicated inhibitor-treated cells had reached a steady value. Vertical axes are arbitrary values, horizontal axes (time) rescaled for uniformity. In each case fluorescence reached a new equilibrium value rapidly after the externally applied shifts in temperature, as indicated (°C). (D) Variation of MTY fluorescence in PBS with temperature. Each data point is the mean ± SD from five independent experiments. The dotted line indicates the slight deviation from linearity at higher temperatures, implying that MTY-based mitochondrial temperature may be over-estimated by ∼2 °C in the 50-60 °C range. (E, F, G) Response of MTY fluorescence to changes in (E) calcium concentration, (F) hydrogen peroxide and (G) pH over time (since first addition of supplement to PBS). Supplements were added sequentially to the cuvette, to bring samples to successively higher concentrations (E, F) or pH (G). Each trace is a mean from 5 independent experiments. Actual pH changes were verified in parallel samples. Note that some traces showed a very slight but continuous bleaching effect with time that was independent of the addition of the supplements and was seen also in control traces for unsupplemented samples analysed in parallel: this has been corrected for, by realigning traces back to the horizontal. (H) Idealized scheme for cellular/mitochondrial fractionation (see Materials and Methods for technical details). Cells are initially lysed by homogenization under hypotonic conditions, and nuclei (blue), cytosol (grey) and membranous debris (black) are removed by a series of low and moderate-speed centrifugation steps, yielding a pellet highly enriched in mitochondria (red). Mitochondria are then resuspended and subfractionated by mild detergent treatment and further centrifugation, separating mitoplasts (red, bounded by inner membrane in black) from the fraction containing the disrupted outer membrane (dark green) and inter-membrane space (IMS) components (light green). In a second subfractionation step, mitoplasts are sonicated and ‘inside-out’ sub-mitochondrial particles (SMPs, black) separated from matrix components (red) by higher speed centrifugation.

**Supplementary Figure S3.**
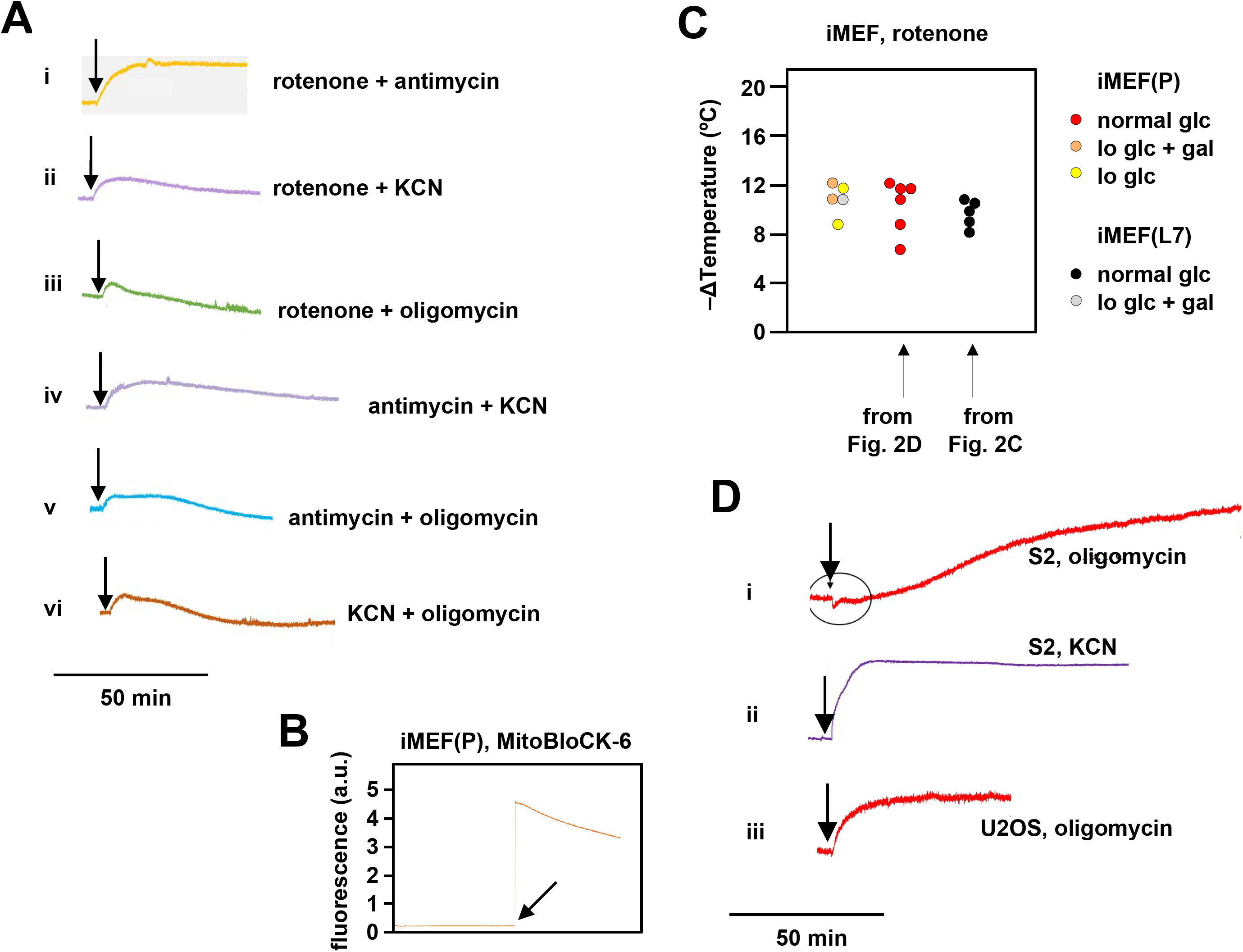
Supplementary data on effects of OXPHOS inhibitors on mitochondrial temperature. (A) Representative fluorescence traces, uniformly rescaled, for MTY-labelled iMEF(P) cells treated with the indicated combinations of OXPHOS inhibitors. Vertical scale (not shown): arbitrary values to similar scale. Rotenone + antimycin (panel i) gave the same stable fluorescence reading as all individual inhibitors, whereas other combinations (panels ii-vi) showed prolonged and/or continuous decline of the fluorescence reading, which is likely be due to dye leakage upon loss of membrane potential, as shown previously^6^ (see Discussion for further exploration of this issue). For this reason we did not attempt to interpret findings from combinations of inhibitors other than rotenone *plus* antimycin (Fig. 2D). (B) Fluorescence trace for addition of 100 μM MitoBloCK-6 to MTY-stained cells. Note the immediate >20-fold increase in fluorescence intensity upon addition of the drug. (C) Extrapolated mitochondrial temperature shifts for iMEFs grown in different media and treated with rotenone. Data for lines iMEF(L7) and iMEF(P) grown in normal glucose (glc) medium are from the experiments shown in Fig. 2C and 2D, respectively. Gal – galactose. Lo glc – low-glucose medium (see Materials and Methods). Data are shown as scatter plots to indicate the distributions. There are no significant differences between the groups (one-way ANOVA with Tukey HSD *post-hoc* test). (C) Representative, uniformly scaled traces of MTY fluorescence in S2 (panel i) and U2OS (panel iii) cells treated with oligomycin and S2 cells treated with KCN (panel ii), as indicated. For clarity, vertical scale (arbitrary values) and temperature calibration are omitted. Fluorescence in oligomycin-treated S2 cells had not reached a stable value even 2 h after addition of the drug; therefore, no meaningful calibration could be conducted. Treatment of other cell-lines with oligomycin or of S2 cells with other OXPHOS inhibitors did not produce this problem, as per the examples shown.

**Supplementary Figure S4.**
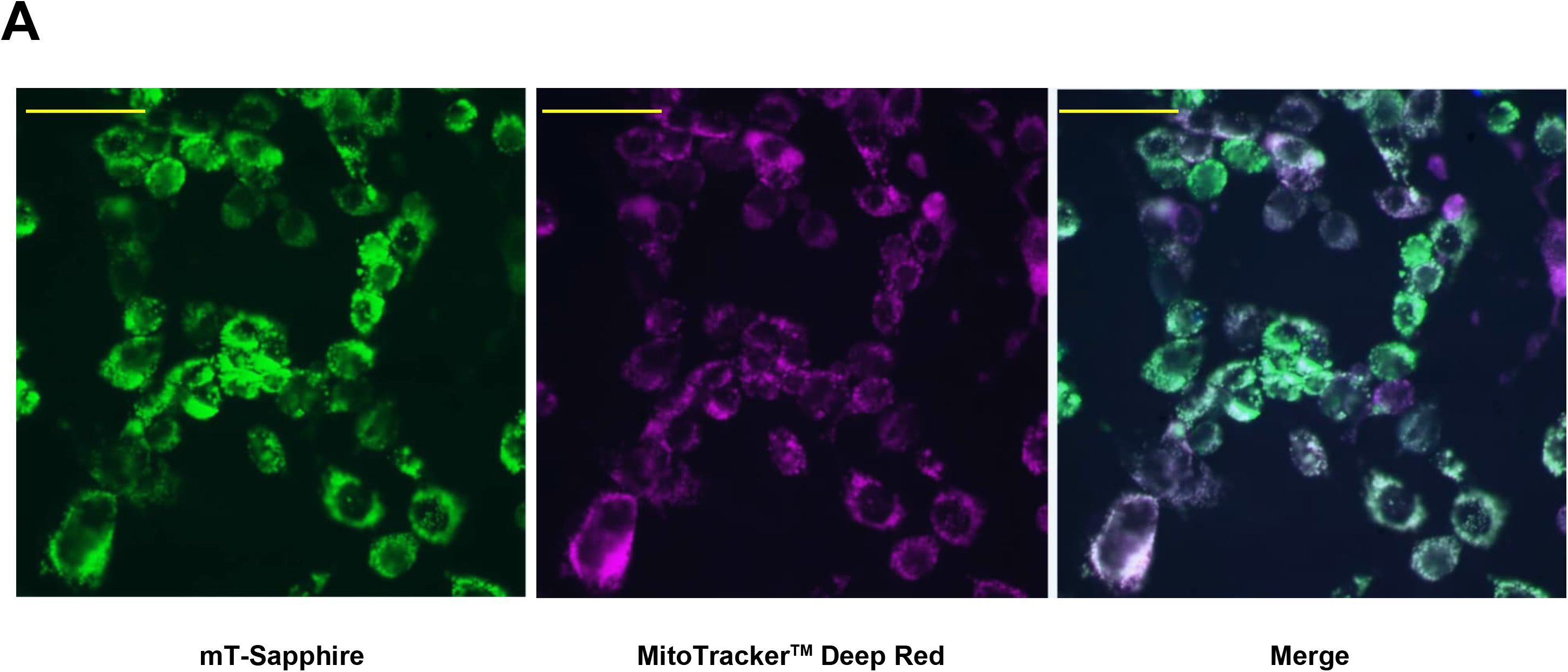
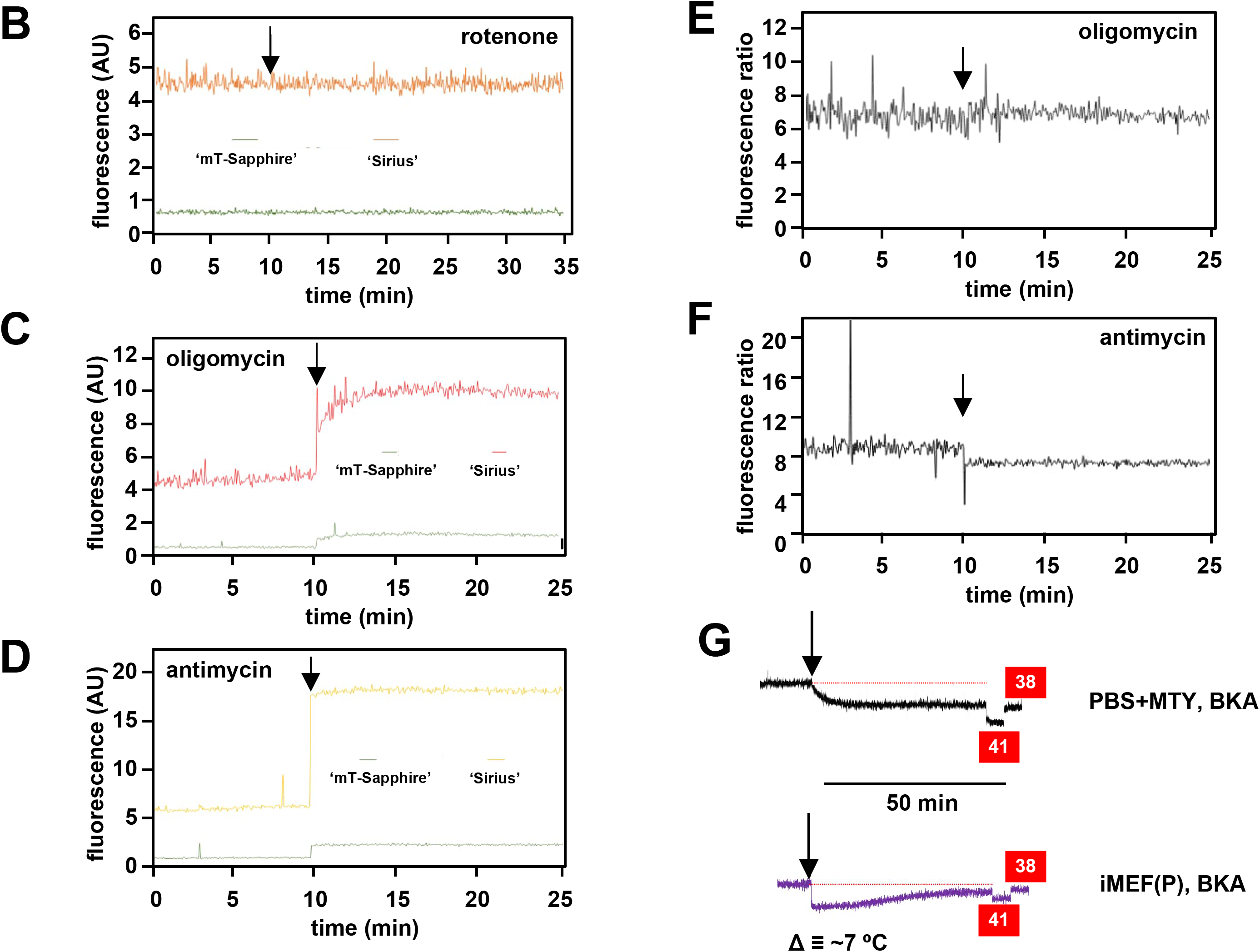
Supplementary data on mitochondrially targeted temperature-sensitive fluorescent probes. (A) Fluorescence microscopy images of iMEF(P) cells transfected with mito-gTEMP and imaged as indicated, as described in Materials and Methods. Scale bars 10 μm. (B, C, D) Fluorescence traces (AU – arbitrary units) and (E, F) fluorescence ratios at the wavelengths used to measure mT-Sapphire and Sirius, in PBS at 38 °C, to which were added the indicated drugs (arrowed). (G) MTY fluorescence traces for cell-free PBS or iMEF(P) cells treated with bongkrekic acid (BKA) as indicated. The drug produced an immediate quenching of fluorescence in both cases. In cells, this was followed by a gradual regain of fluorescence, possibly indicative of mitochondrial temperature decrease by several °C, based on the internal calibration (red boxes). See also Fig. 5E, 5F.

**Supplementary Figure S5.**
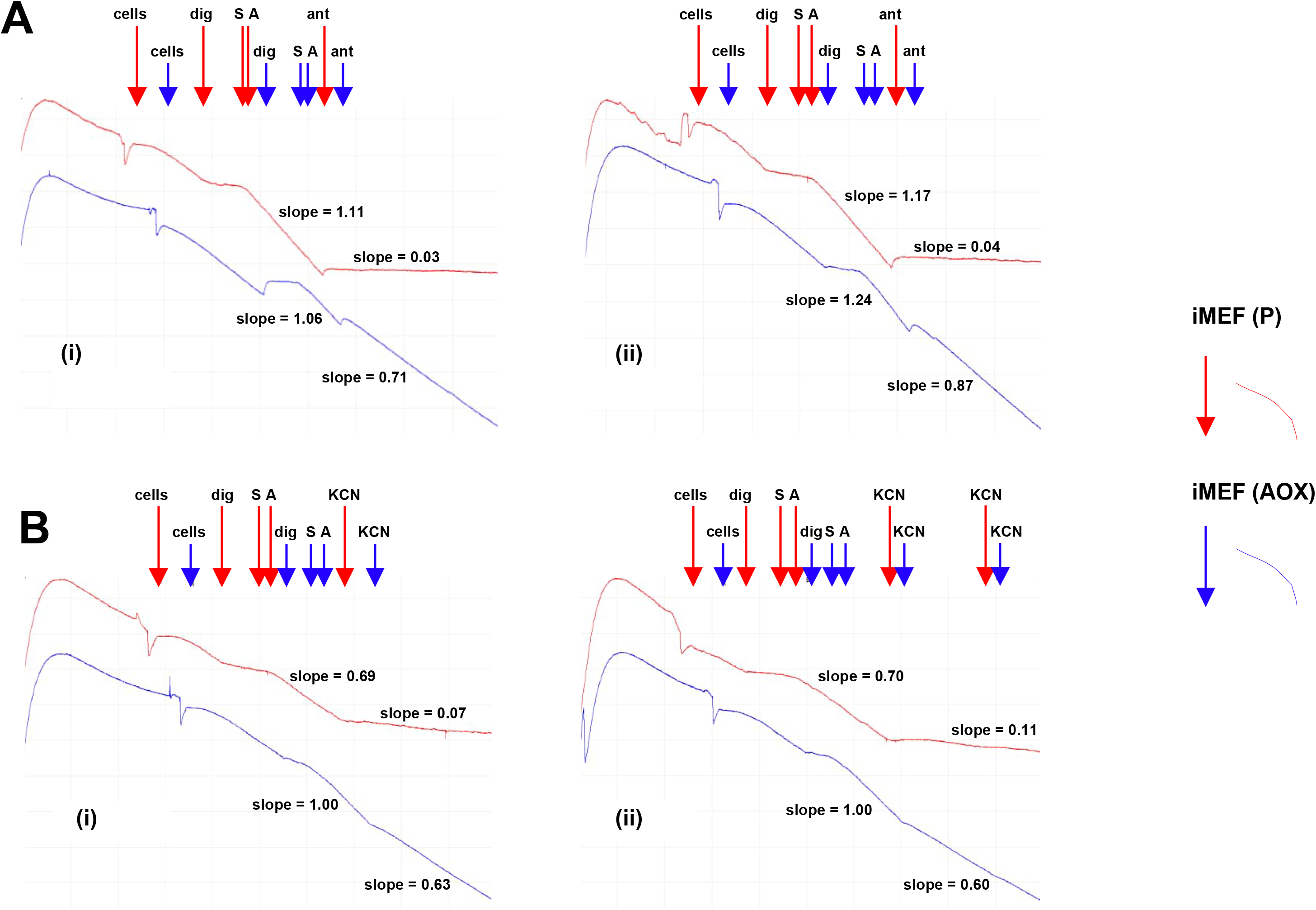
AOX-expressing iMEFs show antimycin- and KCN-resistant respiration. Respirometric traces from digitonin-permeabilized cells of the indicated lines, treated successively with digitonin – dig, pyruvate+glutamate+malate substrate mix – S and ADP – A, followed by either (A) antimycin (ant) or (B) either one or two successive doses of KCN. For details see Methods. In each part, panels (i) and (ii) represent two independent trials. Relative rates of inhibitor-resistant oxygen consumption of iMEF (AOX) cells, based on the indicated slopes, and following subtraction of inhibitor-resistant oxygen consumption of control iMEF (P) cells, were as follows: (A) 70% and 67% antimycin resistance, (B) 50% and 47% KCN resistance.

**Supplementary Figure S6.**
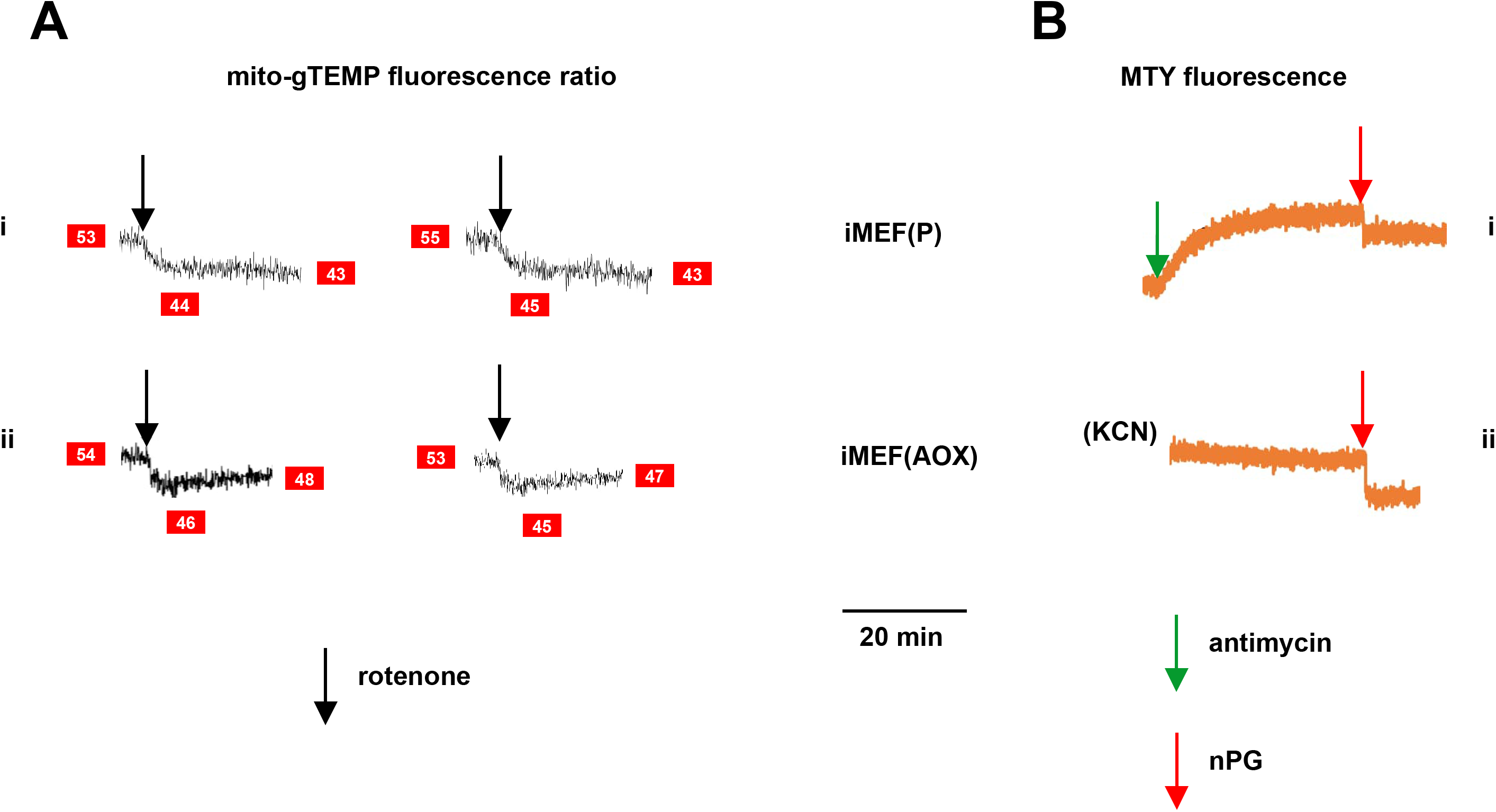
Supplementary data on effects of inhibitors on fluorescence in AOX-expressing cells. (A) Representative traces of mito-gTEMP fluorescence ratio for (I) control iMEF(P) and (ii) iMEF(AOX) cells, after rotenone delivery at 15 min until 35 min, with continuous recording (two replicate experiments in each case). Text in red boxes indicates inferred mitochondrial temperature (to nearest °C) from calibration curve of Fig. 3D. AOX-expressing cells show a consistent warming of ∼2 °C by 20 min post-drug treatment, consistent with the MTY data of Fig. 5C. (B) Representative traces of MTY fluorescence for (i) control iMEF(P) cells and (ii) iMEF(AOX) cells treated with nPG (red arrows). Note that, prior to nPG addition, fluorescence in both cases had reached a stable level – in control cells after antimycin treatment and in AOX-expressing cells after rewarming, following KCN treatment. In both cases, nPG addition produced an abrupt quenching of the fluorescence, making its effect on mitochondrial temperature unquantifiable.

**Supplementary Figure S7.**
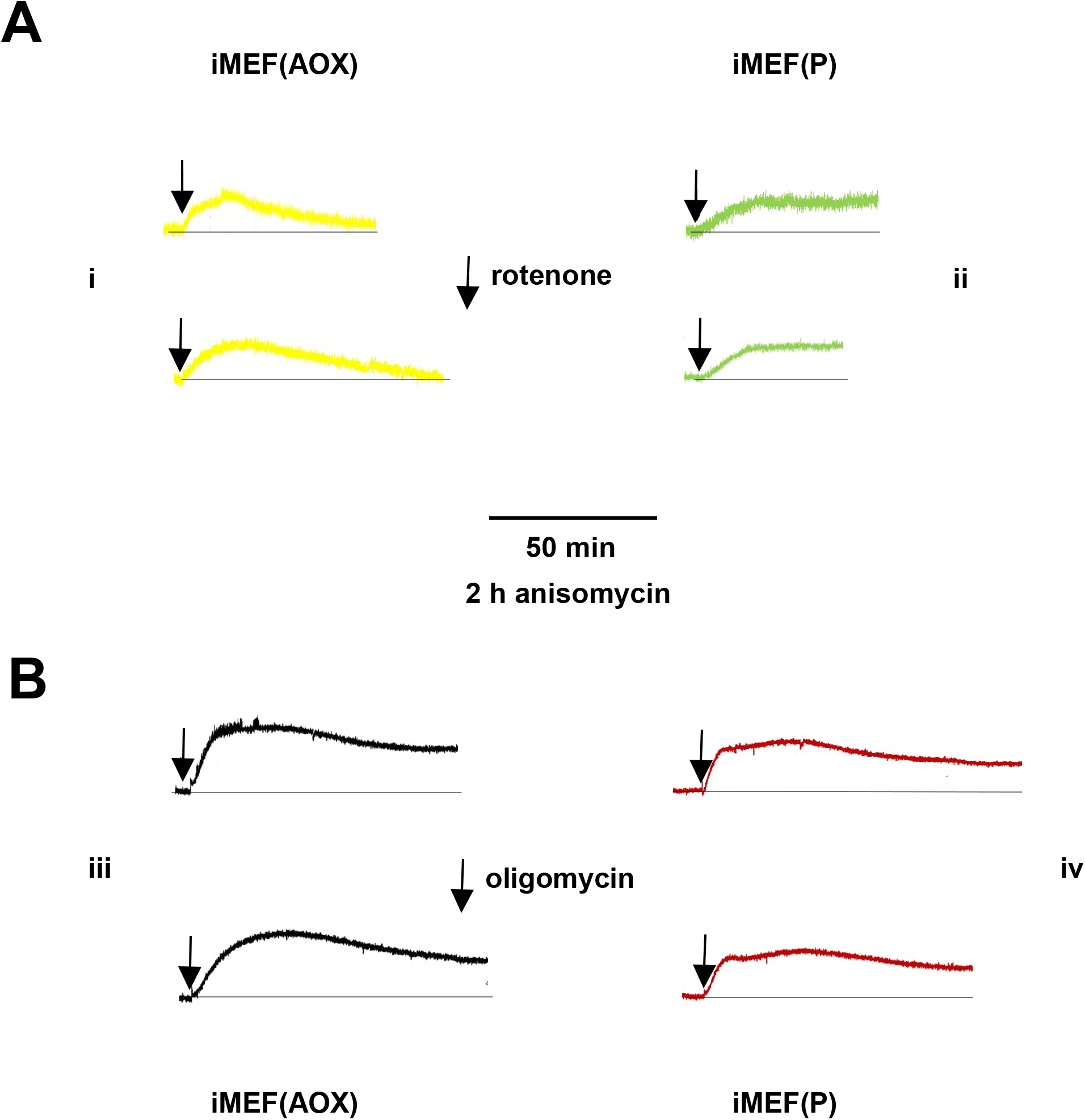
Supplementary data on mitochondrial temperature decrease in anisomycin-treated cells. Representative fluorescence traces, uniformly rescaled, for (panels i, iii) MTY-labelled iMEF(AOX) and (panels ii, iv) iMEF(P)cells pretreated for 2 h with anisomycin, then incubated with (A) rotenone and (B) oligomycin.

## LEGEND TO SUPPLEMENTARY MOVIE

**Supplementary Movie S1**

**Mitochondrial temperature variation between and within cells**

Time-lapse micrographic images (1 frame per 10 s) of iMEF cells stained with MTY, brightness and contrast optimized, showing bright (i.e., potentially cooler) foci within the less intensely stained mitochondrial network. Many of these foci appear mobile within the mitochondrial network, over the 5 min time-scale of the movie. See Fig. 4B for a still image taken from the movie.

